# Inhibition of the IL-17A axis Protects against Immune-related Adverse Events while Supporting Checkpoint Inhibitor Anti-tumor Efficacy

**DOI:** 10.1101/2022.01.19.476844

**Authors:** Melissa G. Lechner, Anushi Y. Patel, Willy Hugo, Trevor E. Angell, Mandy I. Cheng, Marissa S. Pioso, Aline T. Hoang, Natalie Yakobian, Ethan C. McCarthy, Ho-Chung Chen, Eduardo D. Rodriguez, Lily Guo, Michael Astourian, Alexandra Drakaki, Pouyan Famini, Antoni Ribas, Maureen A. Su

## Abstract

Checkpoint inhibitor (ICI) immunotherapy leverages the body’s own immune system to attack cancer cells but leads to unwanted autoimmune side effects in up to 60% of patients. Such immune related adverse events (IrAE) may lead to treatment interruption, permanent organ dysfunction, hospitalization and premature death. Thyroiditis is one of the most common IrAE, but the cause of thyroid IrAE remains unknown. Here we present a novel mouse model in which checkpoint inhibitor therapy leads to multi-organ autoimmune infiltrates and show that activation and infiltration of Type 3 immune cells including IL17A^+^ RORγt^+^ CD4^+^ (T helper 17 or Th17) and gamma delta 17 (γδT17) T cells promote thyroid IrAE development. IL-17A^+^ T cells were similarly found in thyroid specimens from cancer patients treated with ICI who developed thyroid IrAE. Furthermore, antibody-based inhibition of IL-17A, a clinically available therapy, significantly reduced thyroid IrAE development in ICI-treated mice. Finally, combination of IL-17A neutralization with ICI treatment in multiple tumor models did not reduce ICI anti-tumor efficacy. These studies suggest that targeting Th17 and γδ17 function may reduce IrAE without impairing ICI anti-tumor efficacy and may be a generalizable strategy to address IL17-mediated IrAE.

## INTRODUCTION

Immune checkpoint inhibitors (ICI) against programmed death protein (PD-1) and cytotoxic T lymphocyte antigen (CTLA)-4 have revolutionized cancer therapy and hold great promise for the treatment of advanced malignancies^1^. Since initial approval in 2011, eight ICI have entered clinical care, with nearly half of patients with cancer in the U.S. eligible for ICI treatment^2^. ICI are used for first line therapy of metastatic renal, bladder, head and neck, liver, and certain lung, colon, and breast cancers^2^. However, the benefits and use of ICI are limited by development of autoimmune side effects, termed immune related adverse events (IrAEs) seen in up to 60% of patients^3–5^. Endocrine, gastrointestinal and liver, and dermatologic tissues are some of the most common affected by IrAEs^3, 4, 6^. Pulmonary, neurologic, renal, and cardiac autoimmune side effects from ICI are less frequent but associated with grave clinical consequences, including mortality rates of 17-40%^7^. IrAEs can lead to permanent organ dysfunction, cancer therapy interruption, hospitalization, and premature death^5, 7^. With the expanding use of ICIs, IrAEs represent a growing clinical problem.

Despite significant efforts to date, the mechanisms of IrAEs remain poorly understood^8^. Thyroiditis is one of the most common ICI-associated IrAE, occurring in approximately 10-15% of patients treated with anti-PD1/L1 monotherapy and 30% of patients treated with combination anti-PD-1/L1 and CTLA-4^6, 9, 10^. Like IrAEs in other organs, ICI-thyroiditis has both overlapping and distinct features with spontaneous autoimmunity (*i.e.* Hashimoto’s thyroiditis, HT)^9–12^. Two prior important studies^13, 14^ showed intrathyroidal lymphocyte accumulation in ICI-thyroiditis, similar to HT. The onset of thyroiditis during ICI-treatment may be due to the release of immune checkpoints on pre-existing, thyroid-reactive T cells^15^, as suggested by the progressive loss of gland function in patients with HT and increased risk of ICI-thyroiditis in patients with baseline thyroid autoantibodies to thyroid peroxidase (TPO)^16, 17^. On the other hand, thyroid gland destruction in ICI-thyroiditis is more rapid compared to spontaneous HT and often with a clear hyperthyroid phase during gland destruction and subsequent evolution to permanent hypothyroidism^9, 12^. Many patients with ICI-thyroiditis also lack the usual anti-thyroid antibodies to TPO or thyroglobulin (Tg) seen in HT at diagnosis^9, 12^. Thus ICI-thyroiditis, while similar to HT, may be driven by the additional presence of novel immune mechanisms and antigens. Thus, mechanisms underlying ICI-thyroiditis and HT may be shared and/or distinct.

To develop ICI therapies with less toxicity while preserving anti-tumor efficacy, we must determine the cause of IrAE. Currently guidelines recommend suspension immunotherapy and high dose glucocorticoid therapy for severe manifestations (grade 3 or 4) of most IrAE^18, 19^, which may unnecessarily impair the efficacy of ICI cancer treatment for many patients^5, 20^. It is now recognized that high dose glucocorticoid therapy does not attenuate or prevent thyroiditis, hypophysitis, or diabetes, and may correlate with shorter overall survival^12, 21, 22^. Recent data in gut ICI-induced IrAE suggested that immune adverse events may be uncoupled from anti-cancer efficacy using anti-inflammatory treatments^23^. Studies of ICI-associated IrAE in the thyroid may identify common mechanisms driving IrAE development in organs with more grave clinical consequences (*e.g.* pneumonitis). In this study, we define immune mechanisms underlying thyroid IrAE from checkpoint immunotherapy using a newly developed mouse model and translational studies in patients with ICI-thyroiditis. High dimensional analysis of peripheral and thyroid-infiltrating immune cells revealed recruitment and activation of Type 3 immune cells including IL17A^+^ RORγt^+^ CD4^+^ (T helper 17 or Th17) and gamma delta 17 (γδT17) T cells. Th17 cells have previously been implicated in HT^24–26^, suggesting at least partial overlap in pathogenesis between HT and ICI-thyroiditis. Finally, pre-clinical studies demonstrate that IL-17A blockade effectively reduces ICI-thyroiditis without negatively impacting ICI-mediated anti-tumor immunity.

## RESULTS

### Immune infiltrates in thyroid specimens from cancer patients with ICI-thyroiditis

In cancer patients treated with ICI, the thyroid gland is a frequent site of ICI-associated autoimmunity^6^. However, little is known about the immune cell composition in ICI-thyroiditis. To address this, we evaluated thyroid immune cell infiltrates in thyroid FNA biopsy specimens from patients with cancer treated with ICI therapy who developed thyroid inflammation (ICI-thyroiditis, *n*=6) and compared them to subjects with Hashimoto’s thyroiditis (HT, *n*=6), a form of spontaneous thyroid autoimmunity characterized by the formation of B and T cell follicles in the thyroid^27^ (Fig. 1a). Patient demographic and clinical data are shown in **Extended Data Table 1**. As expected from previous reports^9, 10, 28^, patients with HT were predominantly female, whereas the sex distribution was more equal for ICI-thyroiditis. HT patients all had elevated levels of thyroid autoantibodies, whereas this was less common in ICI-thyroiditis patients (one patient had TPO antibodies and two had Tg antibodies). ICI-thyroiditis patients had received combination anti-PD-1 and anti-CTLA-4, anti-PD-1, or anti-PD-L1 therapies for a variety of solid malignancies.

**Figure 1.**
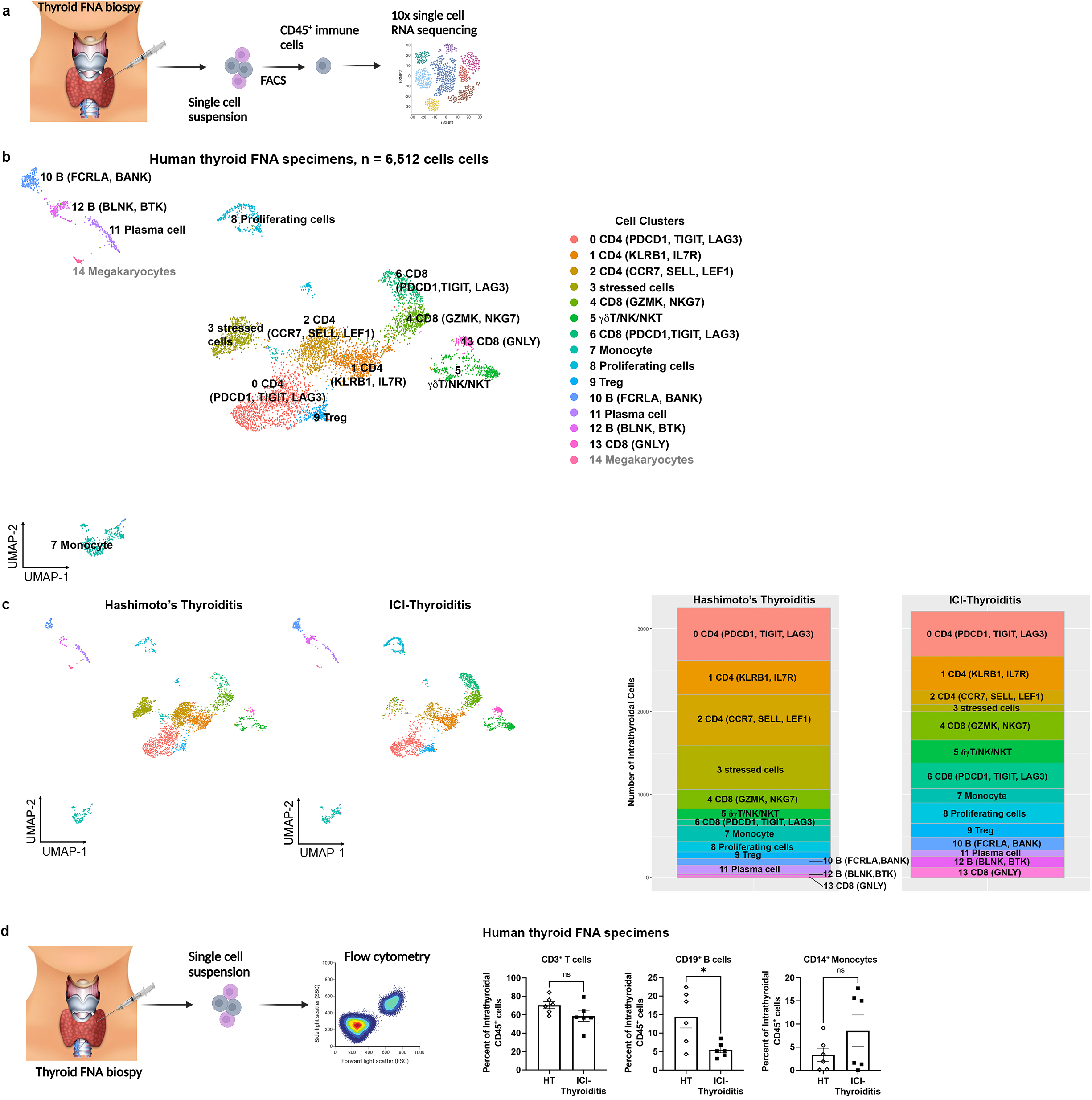
Characterization of thyroid-infiltrating immune cells in ICI-thyroiditis and Hashimoto’s thyroiditis (HT) patients. **a**, Schematic of specimen processing for single cell RNA sequencing of human thyroid fine needle aspiration (FNA) biopsies from patients with Hashimoto’s thyroiditis (HT) or immune checkpoint inhibitor (ICI)-thyroiditis. **b**, scRNAseq data with UMAP plot of 6,512 intrathyroidal CD45^+^ immune cells from patients with HT (*n*=3) or cancer and ICI-thyroiditis (*n*=4). Cluster analysis yields 15 distinct clusters comprising CD4, CD8, and γδ T cells, natural killer (NK)/NKT cells, B cells, monocytes, as well as clusters with mixed cell populations clustered by markers of cell proliferation (proliferating cells) and stressed cells (stressed/dying), and megakaryocytes. **c,** Split plot showing distribution of cells by condition (*left).* Stacked bar plot showing the composition by cell count for pooled specimens in each condition for relevant immune populations; megakaryocytes not shown (*right)*. **d**, Schematic of specimen processing for flow cytometry of human thyroid FNA specimens, and relative frequency of CD3^+^ T cells, CD19^+^ B cells, and CD14^+^ monocytes among CD45^+^ cells in the thyroid FNA specimens from HT or ICI-thyroiditis patients (*n*=6 each) by flow cytometry. Data are mean±SEM. *p<0.05, two-tailed, unpaired t test with Welch correction, assuming unequal s.d.

With the goal of obtaining a broad, unbiased view of the detailed cellular composition associated with ICI-thyroiditis and HT, we turned to single cell RNA sequencing (scRNAseq) of thyroid-infiltrating immune cells from patients with new onset ICI-thyroiditis or HT (Fig. 1c). This technique facilitates identification of rare cell populations and characterization of limited tissue specimens, as in the case of patient thyroid FNA specimens (approximately 10,000 cells per aspirate). Live, single CD45^+^ cells from thyroid FNA specimens from ICI-thyroiditis (*n*=4) or HT (*n*=3) patients were used to construct Chromium™ 10x single cell 5’ gene expression libraries for single cell sequencing. Fifteen heterogeneous immune cell populations were identified and visualized by uniform manifold approximation and projection (UMAP) (Fig. 1c; total *n*=6,512 cells, with 3,292 cells from HT and 3,220 cells from ICI-thyroiditis subjects). Putative cell cluster identities in Fig. 1c were made based upon query of the differentially expressed gene set for each cluster using curated public databases (Enrichr^29^; full gene lists provided in **Extended Data Table 2**). Specifically, lymphoid cell clusters included CD4^+^ T cells (*CD3E, CD4*), CD8^+^ T cells (Cd8: *CD3*, *CD8A, CD8B*), and gamma delta (γδ) T cells (*CD3E, TRDC, variable TRGV genes*). T cell clusters were further defined as CD4 (*PDCD1, TIGIT, LAG3*), CD4 (*KLRB1, IL7R*), CD4 (*SELL, CCR7, LEF1),* CD8 (*GZMK, NKG7*), CD8 (*GNLY*), CD8 (*PDCD1, TIGIT, LAG3*), Treg *(CD4, FOXP3, CTLA4, IKZF2)*. Other immune clusters were monocytic myeloid cells (Mono/Mac: *CD14, FCER1G, CSF1R*), 1B cell (*CD19, CD79A, FCRLA, BANK*), B cell (*CD19, CD79A, BLNK, BTK*), plasma cells (*CD79A*, *IGKC*, various *IGH* and *IGL* genes). Other cell clusters included mixed populations of cells segregated primarily by genes related to cell division (*MKI67, CCNA2, CCNB2*) or consistent with proliferating cell populations (proliferating cells), or genes related to cell stress (stressed: long non-coding RNA, e.g. *NEAT1*). Expression of key genes for each cluster is shown in **Extended Data Fig. 1b.** These scRNAseq data highlight diverse immune cell populations, including multiple populations of activated and exhausted T cells, in thyroid immune infiltrates of ICI-thyroiditis and HT patients.

UMAP projections split by condition demonstrated that cells originating from both HT and ICI-thyroiditis contributed to each cluster (Fig. 2c). The presence of CD4^+^ T cells expressing *PDCD1*, *TIGIT*, and *LAG3* (Cluster 0), and CD4^+^ T cells expressing *KLRB1* and *IL7R* (Cluster 1) was comparable in HT and ICI-thyroiditis specimens (Fig. 2d). In addition, a slight expansion of CD8^+^ T cells expressing exhaustion/checkpoint genes *PDCD1*, *TIGIT*, and *LAG3* (Cluster 6), and cytotoxicity associated genes *GZMK* and *NKG7* (Cluster 4) or *GNLY* (Cluster 13) was seen in ICI-thyroiditis compared to HT (Fig. 2d). Cluster 5, comprised of γδT/NK/NKT cells, was also expanded in ICI-thyroiditis. Finally, ICI-thyroiditis showed fewer plasma cells (Cluster 11), but similar frequencies other B cell populations (Clusters 10 and 12), compared to HT. These data suggest the presence of T, B, and myeloid cell populations in HT and ICI-thyroiditis subjects.

**Figure 2.**
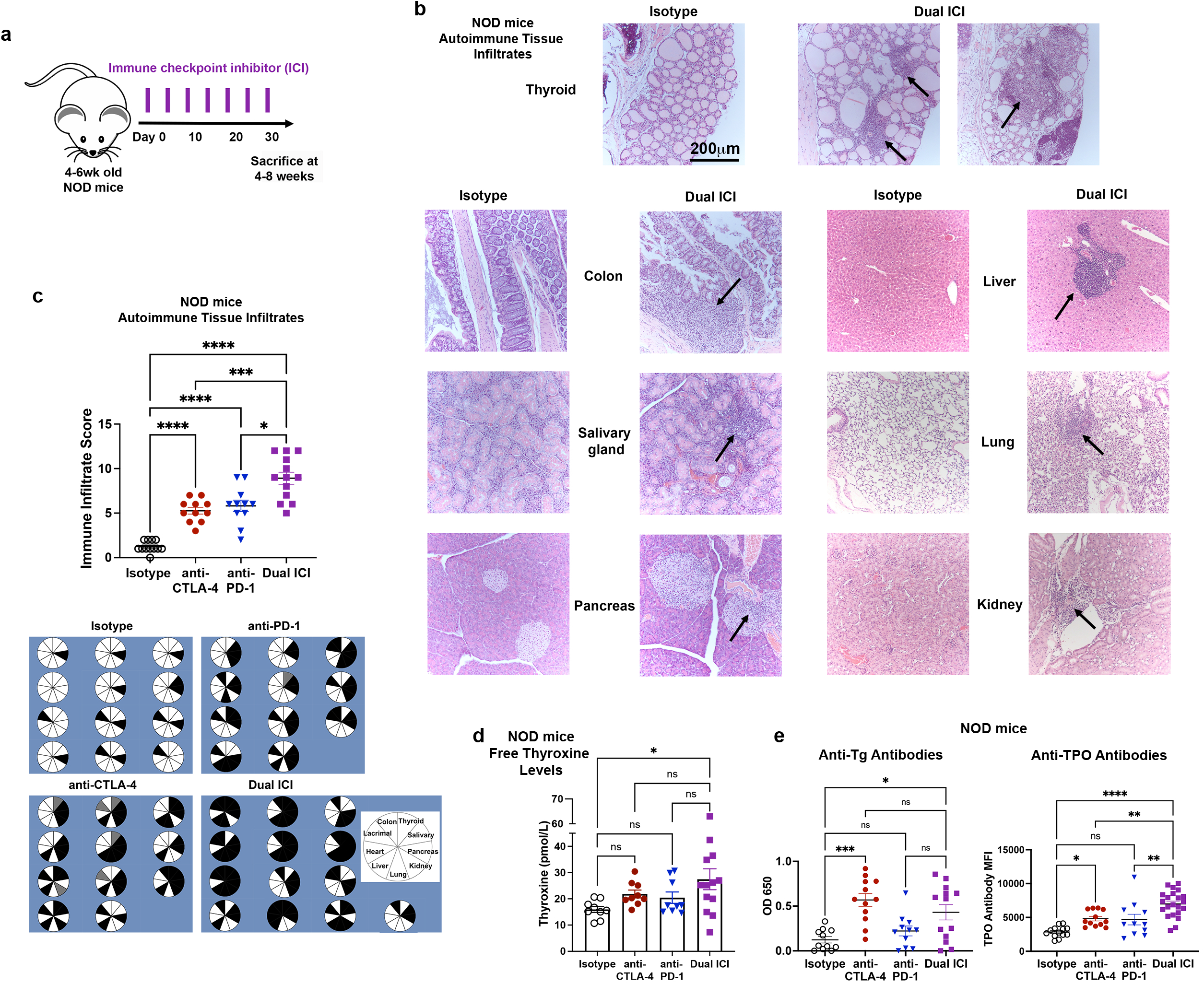
ICI therapy induces multi-system autoimmune infiltrates (IrAE) in NOD mice. **a**, Schematic of ICI drug treatment. **b**, Representative H&E micrographs of autoimmune tissue infiltrates in isotype (*left*) *vs.* Dual ICI (*right*) treated mice (original mag. 400x). **c**, Comparison of autoimmune organ infiltration and severity by immune infiltrate score after 8 weeks of ICI treatment in NOD mice (*n*=12 isotype, n=11 anti-CTLA-4, *n*=11 anti-PD-1, and *n*=13 Dual ICI). Pie charts show tissue infiltrate for each mouse; black = immune infiltrate, white = no infiltrate, gray = no data. **d**, Free thyroxine serum levels in ICI-treated NOD mice after 8 weeks of treatment. **e**, Quantification of anti-thyroid autoantibodies in NOD mice after 8 weeks of treatment (*n*=10 isotype anti-Tg or 13 for anti-TPO, *n*=12 anti-CTLA-4, *n*=11 anti-PD-1, and *n*=13 Dual ICI for anti-Tg or 22 for anti-TPO). Data are mean±SEM. *p<0.05, **p<0.01, ***p<0.001, ****p<0.0001, Brown-Forsythe ANOVA, assuming unequal s.d., followed by Dunnett’s multiple comparisons test (**c, e**), or one-way ANOVA and Dunnett’s multiple comparison test (**d**).

We confirmed the presence of these populations in thyroid FNA specimens by flow cytometry (Fig. 1b, using a gating strategy shown in **Extended Data Fig. 1a)**. T cells and monocytes were present in comparable frequencies (as a percent of CD45^+^ intrathyroidal cells) in ICI-thyroiditis and HT; B cells were less frequent in ICI-thyroiditis (Fig. 2b). Thus, the composition of immune infiltrates in ICI-thyroiditis and HT were at least partially overlapping.

### The NOD background strain predisposes mice to ICI-IrAEs that mimic those seen in humans

Based upon our findings in ICI-thyroiditis patients, we sought to evaluate further the mechanisms underlying development of ICI-associated autoimmunity. The study of ICI-associated IrAE has been hindered by the lack of robust preclinical models^8^. While prior studies have reported autoimmunity in ICI-treated mice, these models on the C57BL/6 (B6) background developed minimal ICI-IrAEs. Augmenting these mild phenotypes required significant manipulation, such as regulatory T cell depletion^30^, genetic knock-in of human checkpoint proteins^31^, chemical stimulants^32^ or co-administration of complete Freund’s adjuvant^20^. A recent paper utilized CBA/J mice in a thyroid IrAE model, but IrAE induction required non-physiologic immunization with human thyroglobulin^33^. We sought to develop a murine model of ICI-IrAE that better recapitulated the development of IrAE in patients; namely, treatment-dependent incident multi-system autoimmune infiltration in an immune competent host.

Underlying subclinical autoimmunity or genetic predisposition increases risk of developing IrAE during ICI cancer therapy in patients^16, 21, 34^. For example, patients with thyroid auto-antibodies against thyroid peroxidase at baseline were five-fold more likely to develop thyroiditis with ICI treatment^11, 35^. The NOD mouse is an autoimmune-prone inbred strain that is best known for its contributions to type 1 diabetes studies but has also been invaluable in understanding generalized autoimmunity^36^. NOD mice develop spontaneous autoimmunity in multiple tissues, including thyroiditis at low frequency (14% at 52 weeks)^37, 38^. Thus, we tested whether ICI treatment in NOD mice could produce robust and reproducible autoimmunity. Groups of four-to six-week old NOD mice were treated twice weekly with anti-mouse CTLA-4 (clone 9D9), anti-mouse PD-1 (RPM1-14), combination (Dual ICI: anti-PD-1 plus anti-CTLA-4), or isotype control antibodies (2A3, MPC-11), at 10 mg/kg/dose *i.p.*, (Fig. 2a). Mice were monitored at least twice weekly for signs of autoimmunity, including weight loss, decreased activity, and glucosuria. As expected, ICI-treated NOD mice showed acceleration of underlying autoimmune risk (*i.e.* diabetes)^11, 21^. Mice developing diabetes were treated with insulin, as described previously^39^. Evaluation of immune cell changes in the spleen occurring with Dual ICI-treatment showed an increased frequency of pan-CD3^+^ T cells, effector/memory (CD44^+^CD62L^-^) subsets among CD4^+^ and CD8^+^ T cells, and regulatory T cells (Treg) compared to isotype-treated control mice (**Extended Data Fig. 2**), consistent with prior reports^40, 41^.

After 4 weeks of treatment, mice were sacrificed, and multiple tissues (salivary, lacrimal, pancreas, liver, lung, heart, colon, eye, gonad, and thyroid) were evaluated by histology and flow cytometry for the development of autoimmune infiltrates. Tissue immune infiltration was quantified by blinded assessment of hematoxylin and eosin (H&E)-stained formalin fixed paraffin-embedded (FFPE) sections (5 high powered fields/section in each animal). Sera were collected for measurement of thyroid autoantibodies to Tg and TPO, as previously described^42^.

ICI-treated NOD mice developed increased immune infiltrates in multiple tissues (*e.g.* thyroid, colon, liver, lung, kidney, and salivary glands) compared to isotype-treated animals, as shown in Fig. 2b and c (ANOVA p<0.0001; p values *vs.* isotype were <0.0001 for anti-CTLA-4, anti-PD-1, and Dual ICI; p=0.0148 for anti-PD-1 and p=0.0008 for anti-CTLA-4 *vs.* Dual ICI, respectively). The predominant inflammation in the thyroid consisted of lymphocytic aggregates within the interstitium and occasionally perivascular aggregates (Fig. 2b). In the liver, both focal collections of lymphocytes within liver parenchyma and peri-portal venule sites were seen. Lung tissues similarly showed aggregates of focal lymphocyte aggregation, as well as loss of alveolar air spaces. Histologic examination of the colon sections showed areas with increased immune infiltrates including basal and intraepithelial lymphocytes. The pattern of immune infiltration in the kidney was primarily glomerulonephritis with accumulation of lymphocytes and peri-vascular lymphocytic aggregates. In the pancreas, immune infiltration was centered on islet and peri-vascular areas, with lymphocytic infiltration of 25-100% of islets in most ICI-treated animals by 4 weeks. Finally, salivary and lacrimal glands showed interstitial lymphocyte aggregates often with diffuse gland involvement. One animal developed pericardial inflammation and one had immune infiltrate in the adrenal gland. Eye, myocardial, and gonad tissues were also evaluated, and autoimmune infiltrates were not seen in any animals. As in patients, autoimmunity occurred more frequently with the combination of anti-PD-1 + anti-CTLA-4 (Dual ICI) *vs.* with single agent ICI (Fig. 2c). Thyroid hormone status was evaluated at 8 weeks by serum measurement of free thyroxine. Dual ICI-treated mice had increased mean thyroxine serum concentrations compared to isotype controls, consistent with excess thyroid hormone release and hyperthyroxinemia during the early destructive phase of ICI-thyroiditis (Fig. 2d). Furthermore, some ICI-treated mice notably exhibited low serum thyroxine consistent with the conversion to hypothyroidism over time in ICI-thyroiditis and mirroring the earlier conversion to hypothyroidism in Dual *vs*. single agent therapy seen in patients.^9^ Some ICI-treated mice developed thyroid autoantibodies. Specifically, anti-TPO antibodies increased with single and dual ICI treatments, and anti-Tg antibodies increased with anti-CTLA4 monotherapy and dual ICI therapy, but not anti-PD-1 (Fig. 2e).

The rate of spontaneous autoimmunity was low in isotype-treated controls which developed only pancreatic insulitis (without diabetes) and occasional lacrimal or salivary gland focal immune infiltration at 12-15 weeks of age (Fig. 2b-c), consistent with their NOD background. Thus, our mouse model recapitulates critical features of clinical ICI-associated autoimmunity, including multi-organ immune infiltration and hyper-and hypothyroidism in a subset of patients, and thus will facilitate studies of the mechanisms leading to IrAEs.

In addition, we evaluated the development of autoimmune infiltrates with ICI treatment in the wild type B6 strain. As done in NOD mice, four-to six-week old B6 mice were treated twice weekly with anti-CTLA-4 and anti-PD-1 antibodies (Dual ICI) or isotype control (**Extended Data Fig. 3a**), and then autoimmune tissue infiltrates and thyroid autoantibodies evaluated. We chose to treat with dual ICI therapy (combination anti-CTLA-4/PD-1) since the combination has been linked with the highest rates of ICI-IrAEs^3, 4^. Overall, B6 mice were resistant to ICI-induced autoimmunity with immune infiltration limited on average to two organs per mouse (**Extended Data Fig. 3b**). Increased immune infiltrates were seen in the liver and lacrimal glands, but not in the thyroid (**Extended Data Fig. 3b-c**). Additionally, B6 mice did not develop thyroid autoantibodies to thyroid peroxidase (TPO) or thyroglobulin (Tg) (**Extended Data Fig. 3d**). Similar results were seen in mice treated with ICI or isotype for eight weeks (data not shown). In summary, B6 mice failed to develop significant multisystem autoimmunity with combination anti-PD1 and anti-CTLA-4 immune checkpoint inhibitor (Dual ICI) therapy. These findings are in keeping with previous reports that B6 mice are resistant to spontaneous autoimmunity, perhaps due to genetic polymorphisms that alter immune responses in this inbred strain^43, 44^.

### Thyroid infiltrating immune cells in ICI-treated mice mirror those in patients with ICI-thyroiditis

ICI-treated NOD mice developed frequent thyroiditis, whereas isotype controls showed low levels of spontaneous autoimmunity consistent with the NOD background strain^36–38^. To delineate immune cell types at single cell resolution in thyroid autoimmune infiltrates in our mouse model, we again turned to scRNAseq (Fig. 3a). Sorted, live CD45^+^ cells from thyroid specimens of NOD mice treated with combination anti-PD-1 and anti-CTLA-4 (Dual ICI, *n*=16 animals) or isotype control (*n*=10 animals), were used to construct Chromium™ 10x single cell 5’ gene expression libraries for single cell sequencing. Seventeen heterogeneous immune cell populations were identified and visualized by UMAP (Fig. 3b; total *n*=10,275 cells, with 8,923 cells from Dual ICI and 1,352 cells from isotype). Putative cell cluster identities in Fig. 3b were assigned using publicly available curated databases (Enrichr^29^ and the Immunological Genome Project database^45^; full gene lists provided in **Extended Data Table 2**). Specifically, lymphoid cell clusters consisted of CD4^+^ T cells (Cd4: *Cd3e, Cd4*), CD8^+^ T cells (Cd8: *Cd3*, *Cd8a*), and gamma delta (γδ) T cells (*Cd3e, Trdc, Trg-V6/V4/V1*). T cell clusters were further defined as CD4 (*Sell, Ccr7;* and *Lef1, Sell, Ccr7*), Treg *(Cd3e*, *Cd4, Foxp3, Ctla4, Ikzf2),* CD4 and CD8 (*Cd4 or CD8a, Icos, Gzmk*), and CD8 (*Cd8a, Cd8b1, Sell, Ccr7*). γδT cells were found in two clusters, (Cluster 5: *Rorc*^+^ *Tcrg-V6*^+^ and *Trcg-V4*^+^ γδT, and Cluster 7: *Tbx21^+^ Tcrg-V1*^+^ and *Trcg-V4*^+^ γδT). Cluster 7 also included natural killer (NK) and NKT cells (*Ncr1, Cd7, Nkg7, Klrd1, Tbx21*). Other immune cell clusters included macrophages and monocytes (*Cd68, Cd14, Fcgr1, Adgre1, C1qa/b/c, Csfr1, Cd74*), conventional dendritic cells (cDC: *Cd14, Itgax, H2-Aa*), plasmacytoid dendritic cells (pDC: *Siglech, Itgax, Cd14*), innate lymphoid cells type 2 (ILC2: *Il7r, Il1rb, Il13, Il5, Fosb*), and B cells (*Cd79a, Cd19, Cd74, Igkc*). Other cell clusters included two comprised of mixed immune cell types segregated primarily by *Mik67* and *Ccna2*, consistent with proliferating cell populations (4 and 10), stressed or dying cells (14, expression of long non-coding RNA *Malat1*), and two clusters of cervical thymic cells (15 and 16, identity determined by Enrichr and Immunologic Genome Project). Expression of key immune cell genes for each cluster is shown in **Extended Data Fig. 4a.** UMAP projections split by condition demonstrated that cells originating from either isotype-treated or dual ICI-treated thyroids were present in each cluster (Fig. 3b). Notably, numbers of immune cells were higher in ICI-treated thyroids compared to isotype-treated (Fig. 3c). Thus, while the composition of immune cells in ICI-and isotype-treated thyroids were similar, these findings suggest that immune cell populations are highly expanded with ICI treatment.

**Figure 3.**
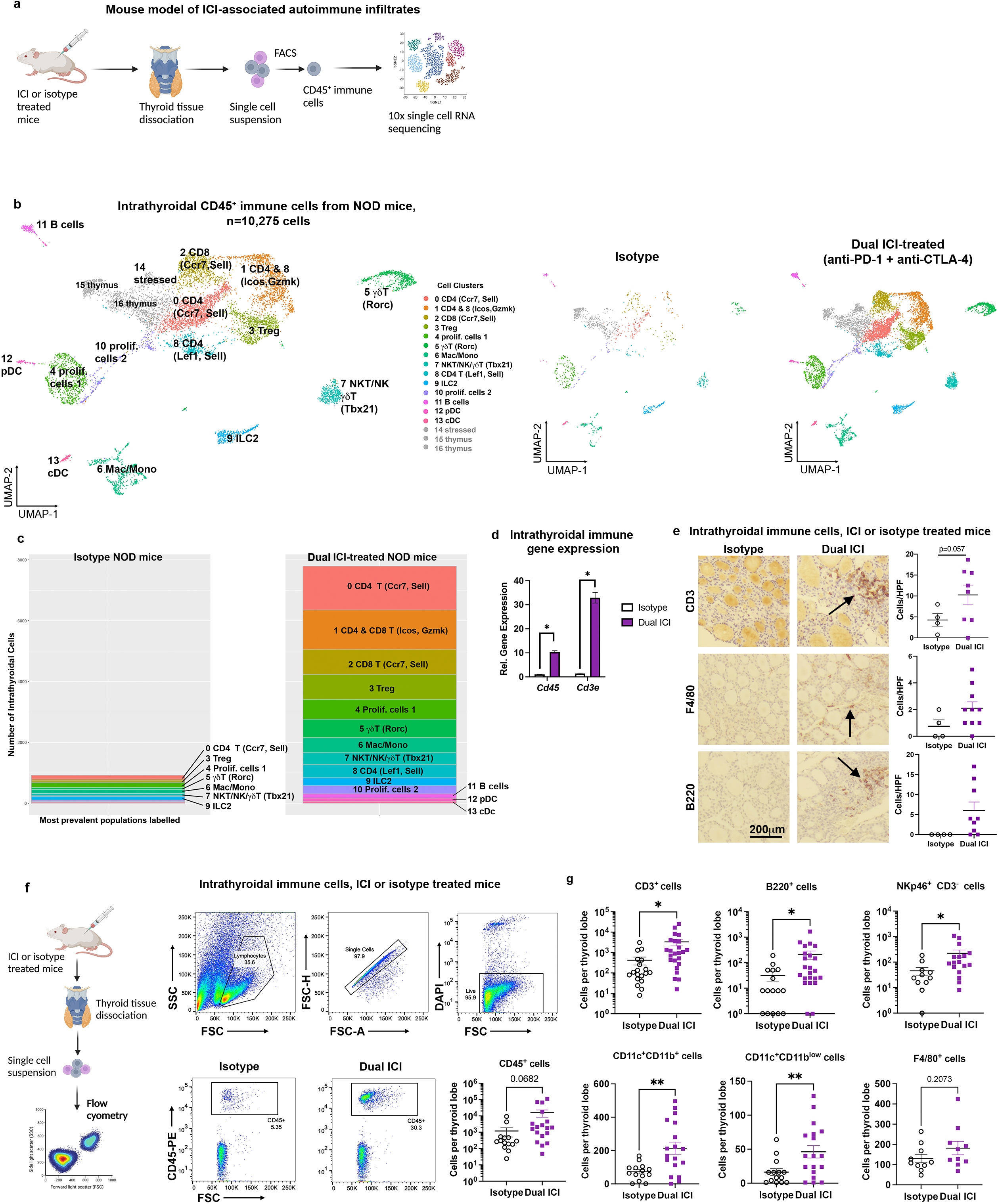
Characterization of thyroid-infiltrating immune cells in ICI-treated NOD mice. **a,** Schematic of thyroid tissue processing for single cell RNA sequencing from Dual ICI or isotype-treated mice. **b,** UMAP plot of 10,275 thyroid-infiltrating CD45^+^ immune cells from Dual ICI-treated (*n*=16) or isotype-treated (*n*=10) mice. Cluster analysis yields 17 distinct clusters comprising CD4, CD8, and γδ T cells, NK/NKT cells, innate lymphoid cells (ILC), B cells, macrophages/monocytes (Mac/Mono), conventional (cDC) or plasmacytoid (pDC) dendritic (DC), as well as clusters with mixed cell populations clustered by markers of cell proliferation (proliferating cells 1 and 2), stressed or dying cells (stressed), or cervical thymic contamination (thymus). **c,** Stacked bar plot showing the composition by cell count for pooled specimens in each condition for relevant immune populations (*left)*. Split plot showing distribution of cells by condition (*right).* **d**, Comparison of intrathyroidal gene expression in isotype *vs.* Dual ICI-treated mice for *Cd45* and *Cd3e*, measured by qRT-PCR. *n*=12 animals/group; thyroid tissue pooled and run in 3 experiments, in triplicate**. e**, IHC staining for T cells (CD3), macrophages (F4/80), and B cells (B220) in thyroid tissue from isotype (n*=4*) or Dual ICI-(*n=*7,10, or 9) treated mice [*left,* representative sections, 400x original mag.; *right*, cells/hpf]. **f-g**, Comparison of thyroid immune cell infiltrate by flow cytometry. Representative gating strategies and dot plots and comparison of accumulated CD45^+^ immune cells (**f**) and subpopulations (**g**) in thyroid tissue from isotype (*n*=11, except *n*=16 for CD3^+^ and B220^+^) *vs*. Dual ICI-treated (*n*=17, except *n*=23 for CD3^+^ and B220^+^) mice after 4 weeks. Data shown as cells per thyroid lobe. Data are mean±SEM. *p<0.05, **p<0.01, two-tailed, unpaired t test with Welch correction, assuming unequal s.d. and Holm-Sidak method correction for multiple comparisons (**e-g**).

Thyroid infiltrating T cells showed gene expression associated with activation (*Cd44, Cd69, Icos*) and effector functions, including cytotoxicity (e.g. *Gzmk)*, as shown in **Extended Data Fig. 4a**. This accumulation of CD4 and CD8 T cells with cytotoxic and effector gene expression mirrored that seen in ICI-thyroiditis and HT patients. Furthermore, T cell checkpoint and exhaustion genes *Pdcd*1 and *Tigit* were also seen in thyroid immune infiltrates of mice, with expression highest in i) γδT cells (Clusters 5 and 7), ii) a mixed CD4 and CD8 population of proliferating T cells (Cluster 10), iii) Treg cells (Cluster 3), and iv) pDC (Cluster 12)^46^ (**Extended Data Fig. 4a)**. Compared to the immune composition of human ICI-thyroiditis specimens, ICI-treated mice similarly showed a predominant accumulation of T cells, including CD4, CD8, and γδT subsets (Fig. 3b and c). Myeloid cells (monocytes, macrophages, and dendritic cells) and B cells were present as minor populations in thyroid infiltrates (Fig. 3b and c), as was seen in ICI-thyroiditis patients (Fig. 1).

Our scRNAseq data showed accumulation of multiple T cell types in the thyroid tissue of ICI-treated mice, as well as in the sparse infiltrates of isotype-treated NOD mice, which have a genetic predisposition to autoimmunity. We used quantitative reverse transcriptase polymerase chain reaction (qRT-PCR) to compare relative intrathyroidal gene expression of *Cd45,* a marker of hematopoetically derived cells, and *Cd3e,* a T cell lineage marker, in ICI-treated *vs.* isotype mice (Fig. 3d). Expression of these genes were significantly increased, consistent with increased thyroid immune cell infiltration, notably T cells, with ICI treatment. We then evaluated the expression of immune cell markers in thyroid infiltrates in ICI-vs. isotype treated NOD mice using immunohistochemistry Fig. 3e. Immunohistochemical staining for canonical immune markers on FFPE sections showed the presence of T cells (CD3), macrophages (F4/80), and B cells (B220) in immune aggregates in ICI-treated mice, with rare immune cells in the thyroid parenchyma of isotype controls (Fig. 3e). To quantify the types of immune cells in thyroid tissue of ICI-vs. isotype treated NOD mice, we used flow cytometry, as shown in Fig. 3f (gating strategy in **Extended Data Fig. 5a**). Fresh thyroid tissues from ICI or isotype-treated mice were perfused with saline to remove circulating peripheral immune cells and dissociated into single cell suspensions then stained for immune markers and analyzed for CD45^+^-gated immune cells. More infiltrating CD45^+^ immune cells were seen in thyroid tissue of ICI-treated mice compared to isotype controls (mean 15,875 cells/thyroid ± SEM 7,515 *vs*. 1,195±687, trend, p=0.068) (Fig. 3f). T cells (CD3^+^) constituted the major group of the infiltrating immune cells (CD45^+^) in thyroid tissue and were significantly increased in ICI-treated mice (mean 3,332±1,220 cells/thyroid lobe *vs*. 420±176, p=0.027) (Fig. 3g). Other populations seen in thyroid immune infiltrates were also more numerous in ICI-treated mice, including DCs (CD11c^+^ CD3^-^ F4/80^-^ B220^-^ CD11b^low^, mean 46±9 *vs.* 16±5, p=0.008; CD11c^+^ CD3^-^ F4/80^-^ B220^-^ CD11b^+^ mean 214±37 *vs.* 73±13, p=0.001), B cells (B220^+^ CD3^-^ CD11b^-^, mean 212±77 *vs*. 32±13, p=0.03), and NK cells (NKp46^+^CD3^-^, mean 221±74 *vs*. 46±16, p=0.03) (Fig. 3g). While macrophages were present in thyroid immune infiltrates, we did not find a significant difference in mean intrathyroidal macrophage number between ICI-and isotype-treated mice (F4/80^+^, mean 182±33 *vs.* 130±21, p=0.21) (Fig. 3g). These data confirm and provide quantification of the thyroid-infiltrating immune cell populations in ICI-vs. isotype-treated NOD mice seen with scRNAseq.

### Evidence for a role for Type 3 immunity in ICI-mediated thyroiditis

Interestingly, scRNAseq data from patient specimens with ICI-thyroiditis showed the presence of rare *RORC*^+^ and *IL23R*^+^ T cells (Fig. 4a). *RORC*^+^*IL23R*^+^ T cell are classically associated with spontaneous thyroid autoimmunity (*e.g.* HT), and indeed were also seen in scRNAseq data of HT patient specimens (Fig. 4a)^24–26, 47^. However, *RORC*^+^*IL23R*^+^ T cells and the role of Type 3 immunity have not yet been described in ICI-thyroiditis. *RORC* encodes RORγt, a master transcriptional regulator that drives expression of IL-17 cytokines in multiple cell types^48, 49^. *IL23R* encodes the receptor for IL-23, a cytokine that stimulates T cell IL-17 production and has been linked with multiple autoimmune diseases, including colitis, psoriasis, and inflammatory arthritis^50^. Therefore, we reasoned that Type 3 immune responses might also underlie autoimmune pathogenesis in ICI-thyroiditis. Type 3 immunity genes, including *Rorc* and *Il23r*, were also seen in scRNAseq of intrathyroidal immune cells from ICI and isotype-treated NOD mice (Fig. 4b), with highest expression in Cluster 5 (**Extended Data Fig. 4b**). Cells in this *Rorc^+^ Il23r* cluster also expressed *Cd3e, Il17a*, *Trdc,* and *Tcrg-V6,* supporting their inclusion of IL-17-producing gamma delta T cells (γδT17) (**Extended Data Fig. 4b**)^51, 52^. The presence of *Rorc^+^ Il23r^+^* T cells in isotype treated mice (Fig. 4b) may reflect low levels of spontaneous thyroid autoimmunity previously described in NOD mice^36, 37^. Thus, our scRNAseq data from both humans and mice, suggest a potential role for RORγt^+^ IL-17A-producing T cells in not only spontaneous thyroiditis, as has been previously reported^24, 26^, but also ICI-associated thyroiditis.

**Figure 4.**
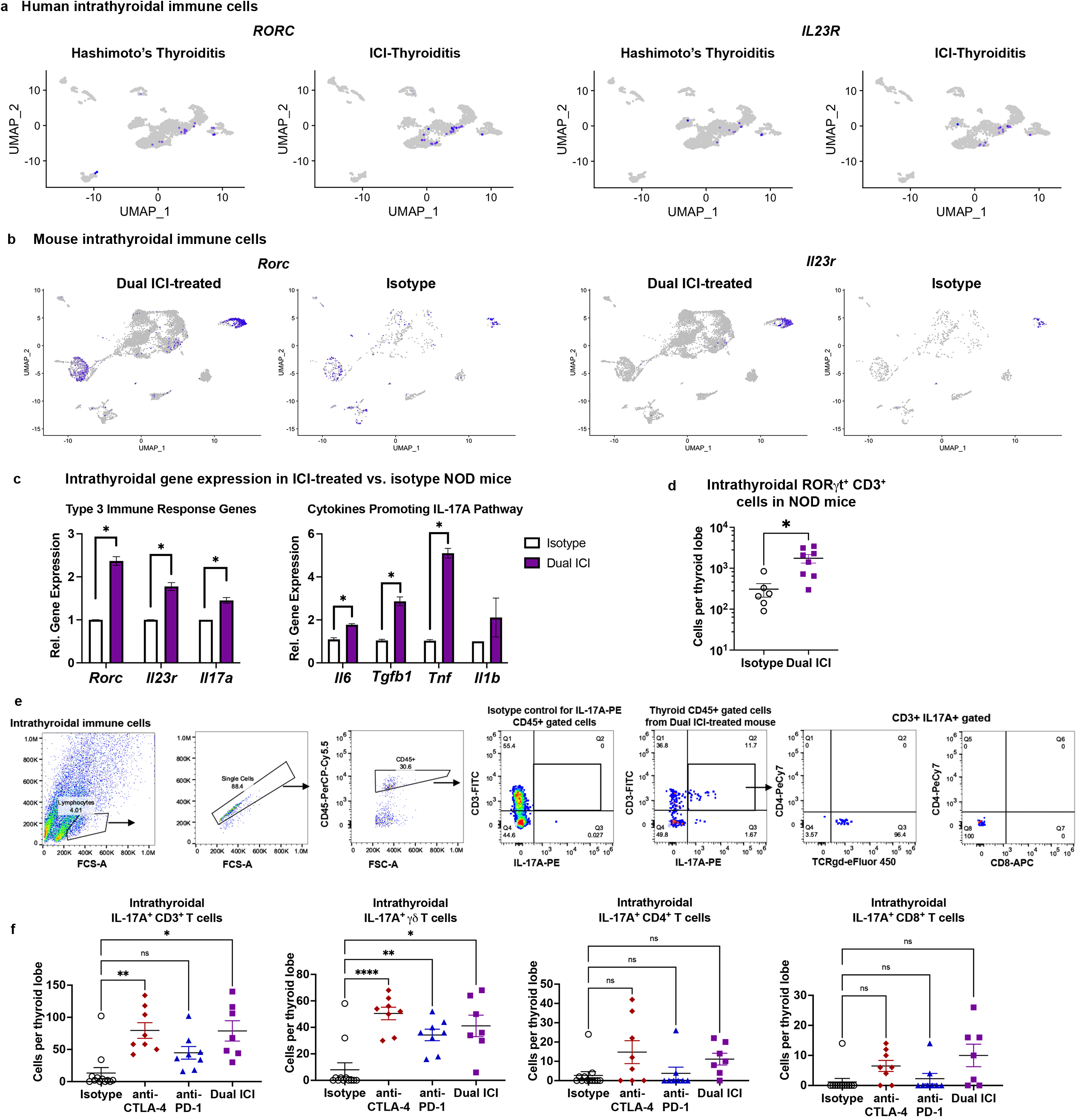
RORγt^+^ IL-17A^+^ T cells in thyroid immune infiltrates of ICI-thyroiditis patients and Dual ICI-treated mice. **a**, Feature plots showing *RORC^+^* and *IL23R^+^* cells associated with T cell clusters in thyroid FNA specimens from patients with Hashimoto’s thyroiditis (*left*) and ICI-thyroiditis (*right*) by scRNAseq. **b**, Feature plots showing *Rorc^+^* and *Il23r^+^* cells associated with T cell clusters in intrathyroidal immune cells from isotype or ICI-treated NOD mice by scRNAseq **c,** Comparison of intrathyroidal gene expression in isotype *vs.* Dual ICI-treated mice for Type 3 immunity (RORγt-pathway) associated genes *Rorc, Il23r, Il17a,* and cytokines associated with IL-17A pathway differentiation (*IL6, Tgfb1, Tnf, Il1b)* measured by qRT-PCR. *n*=12 animals/group; thyroid tissue pooled and run in 3 experiments, in triplicate**. d**, Comparison of thyroid-infiltrating RORγt^+^ pan-CD3^+^ (αβ and γδ) T cells by flow cytometry in isotype (*n*=6) *vs.* Dual ICI-treated (*n*=8) mice. **e**, Representative flow cytometry plots and gating strategy and **f,** quantification of intrathyroidal IL-17A^+^ CD3^+^ T cells and IL-17A^+^ T cell subsets from ICI-treated mice at 4 weeks; isotype (*n*=12), anti-PD-1 (*n*=8), anti-CTLA-4 (*n*=8), and Dual ICI (*n*=7). Data are mean ±SEM shown. *p<0.05, **p<0.01, ***p<0.001, ****p<0.0001. two-tailed, unpaired t test with Welch correction, assuming unequal s.d. (**c, d**) and Holm-Sidak method correction for multiple comparisons (**c**); Brown-Forsythe ANOVA, assuming unequal s.d., followed by Dunnett’s multiple comparisons test (**f**).

We turned to additional modalities to quantify changes in intrathyroidal *RORC*^+^*IL23R*^+^ T cell populations occurring with ICI treatment in our mouse model. First, we confirmed increased expression of *Rorc* and associated Th17/Tc17/γδT17 pathway genes in whole, saline-perfused thyroid tissue from Dual ICI-treated compared to isotype-treated NOD mice using qRT-PCR. As shown in Fig. 4c, expression was significantly increased in ICI *vs.* isotype-treated mice for *Rorc* [mean fold change (FC) 2.4±0.1, p<0.001], *Il23r* (FC 1.28±0.09, p<0.001), and *Il17a* (FC 1.45±0.07, p<0.001)]. Cytokines previously shown to promote IL-17-producing CD4^+^ T helper (Th17)^47^ differentiation (TGFβ and IL-6) and γδT17 activation (TNFα and IL-1β)^52^ were also increased in thyroids of ICI-treated mice compared to isotype-treated animals (Fig. 4c). In addition, flow cytometry analysis of infiltrating immune cells dissociated from fresh thyroid showed increased intrathyroidal RORγt^+^ pan-CD3^+^ T cells [inclusive of CD4^+^ Th17^47^, CD8^+^ T effector (Tc17)^53^, and γδT17^52^ subsets], in Dual ICI-treated mice compared to isotype controls (mean 936±267 *vs.* 183±43 cells per thyroid lobe, p=0.010, Fig. 4d). To better characterize RORγt^+^ and IL-17A-producing populations in thyroid immune infiltrates of ICI-treated mice, we used flow cytometry to quantify different IL-17A^+^ cell subsets: CD4^+^ Th17, CD8^+^ Tc17^53^, and γδT17 cells (Fig. 4e, with additional gating strategy shown in **Extended Data Fig. 5b**). CD3^+^ IL-17A^+^ cells were increased in the thyroids of mice ICI-treated mice (Fig. 4f, mean 13± SEM 8 cells per thyroid lobe in isotype-treated *vs.* 79±12 in anti-CTLA-4 *vs.* 45±10 in anti-PD-1, 79±15 in Dual ICI; ANOVA p<0.001; p values for comparison to isotype were anti-CTLA-4 0.002, anti-PD-1 0.07, Dual ICI 0.01). Among IL-17A^+^ T cell subsets, γδT17 cells were significantly increased in thyroid immune infiltrates following ICI treatment (8±5 cells per thyroid lobe in isotype-treated *vs.* 50±5 in anti-CTLA-4 *vs.* 34±4 in anti-PD-1, 41±8 in Dual ICI; ANOVA p<0.0001; p values for comparison to isotype were anti-CTLA-4 <0.001, anti-PD-1 0.003, Dual ICI 0.02). Th17 and Tc17 cells were not significantly different in ICI-treated groups compared to isotype controls (Fig. 4f). Rare CD3^-^ IL-17A^+^ cells were seen in intrathyroidal immune infiltrates by flow cytometry, which may represent macrophages or innate lymphoid cells type 3 (ILC3)^54^. These data mirror the scRNAseq data above showing *Rorc* expression localized to T cell clusters. Together these data strongly implicate a role for IL-17A^+^ RORγt^+^ T cells, including γδT17 cells, in the development of ICI-thyroiditis.

### IL-17A neutralizing antibody reduces autoimmune infiltrates in ICI-treated mice

Based on these data, we predicted that blocking IL-17A could reduce thyroid autoimmunity in ICI-treated mice. We therefore tested the effects of an IL-17A neutralizing antibody (clone 17F3, 0.5mg/dose 3x/week *i.p.*) 10 days after Dual ICI therapy in NOD mice (Fig. 5a). Indeed, this IL-17A neutralizing antibody reduced the frequency and severity of autoimmune infiltrates in ICI-treated mice at 4 weeks, including thyroiditis (Fig. 5b; ANOVA p<0.0001 for immune infiltrate; p values *vs.* isotype were Dual ICI <0.0001 and Dual + αIL17A 0.005; p value Dual ICI vs Dual + αIL17A 0.003). Specifically for thyroiditis (Fig. 5c, gating strategy shown in **Extended Data Fig. 5**), thyroid infiltrating CD45^+^ immune cells were reduced from mean 16,251± SEM 6,068 cells/lobe in Dual ICI-treated mice, to 339±133 in Dual ICI with anti-IL17A, and compared to 3,389±2,273 in isotype-treated mice (ANOVA p<0.03; p values *vs.* isotype were Dual ICI 0.18 and Dual + αIL17A 0.35; p value Dual ICI vs Dual + αIL17A 0.045). Thyroid-infiltrating CD3^+^(αβ and γδ TCR) T cells were reduced as well, mean 4,031±1,167 cells/thyroid lobe in Dual ICI *vs.* 164±59 with Dual ICI + anti-IL17A *vs.* 651±229 with isotype (ANOVA p=0.001; p values *vs.* isotype were Dual ICI 0.02 and Dual + αIL17A 0.0.14; p value Dual ICI vs Dual + αIL17A 0.008). Other thyroid-infiltrating immune cell populations evaluated were also decreased, including B cells (mean 212±77 cells/thyroid lobe in Dual ICI *vs.* 20±10 in Dual ICI + anti-IL17A *vs.* 32±13 in isotype, ANOVA p=0.01; p values *vs.* isotype were Dual ICI 0.09 and Dual + αIL17A 0.86; p value Dual ICI vs Dual + αIL17A 0.06), NK cells (mean 222±75 cells/thyroid lobe in Dual ICI *vs.* 16±4 in Dual ICI + anti-IL17A *vs.* 46±16 in isotype, ANOVA p=0.009; p values *vs.* isotype were Dual ICI 0.09 and Dual + αIL17A 0.27; p value Dual ICI vs Dual + αIL17A 0.04), CD11c^+^CD11b^low^ DC (mean 42±9 cells/thyroid lobe in Dual ICI *vs.* 8±2 in Dual ICI + anti-IL17A *vs.* 11±3 in isotype, ANOVA p<0.001; p values *vs.* isotype were Dual ICI 0.008 and Dual + αIL17A 0.92; p value Dual ICI vs Dual + αIL17A 0.005), and CD11c^+^CD11b^+^ DC (mean 214±39 cells/thyroid lobe in Dual ICI *vs.* 46±10 in Dual ICI + anti-IL17A *vs.* 67±13 in isotype, ANOVA p<0.001; p values *vs.* isotype were Dual ICI 0.006 and Dual + αIL17A 0.50; p value Dual ICI vs Dual + αIL17A 0.002). Controls treated with isotype + IL17A inhibitor showed infiltrates comparable to isotype-only controls (data not shown). Thus, inhibition of the IL-17A axis (via IL-17A) effectively decreases autoimmune tissue infiltration during ICI treatment.

**Figure 5.**
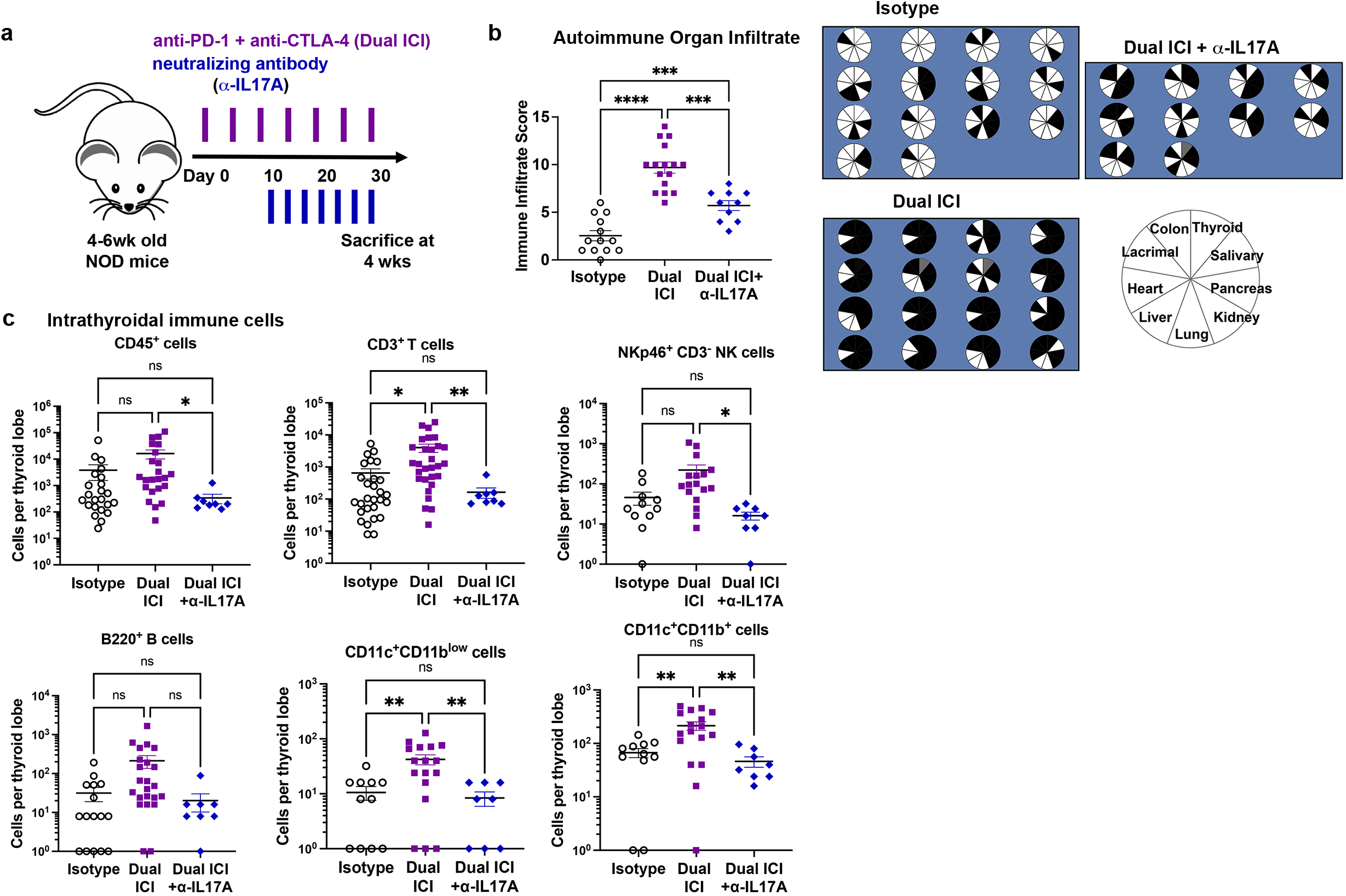
Neutralizing interleukin (IL)-17A antibody therapy reduces ICI-associated autoimmune infiltrates. **a**, Schematic of treatment regimen combining anti-PD-1 + anti-CTLA-4 (Dual ICI) with a neutralizing IL-17A antibody in NOD mice. **b**, Autoimmune organ infiltrate score after 4 weeks of ICI treatment in isotype (*n*=14), Dual ICI (*n*=16), or Dual ICI + neutralizing IL-17A antibody (αIL17A, *n*=10). Pie charts (*right*) showing tissues with immune infiltration after 4 weeks of ICI treatment. Each pie represents one animal; black = immune infiltrate, white = no infiltrate, gray = no data. **c**, Intrathyroidal immune cell frequency among groups [isotype, *n*=11, except *n*=16 for CD3^+^), Dual ICI (*n*=17, except *n*=23 for CD3^+^), Dual ICI + αIL17A (*n*=8)]. Data are mean ±SEM shown. *p<0.05, **p<0.01, ***p<0.001, ****p<0.0001. Brown-Forsythe ANOVA, assuming unequal s.d., followed by Dunnett’s multiple comparisons test (**b, c**).

### Evaluation of ICI-associated thyroid autoimmunity in tumor-bearing NOD mice

We next evaluated thyroid autoimmunity in tumor-bearing NOD mice to test whether the observed increase in IL-17A^+^ T cells with ICI treatment were also seen in the context of a cancer. Few syngeneic tumor models have been developed on the NOD background^55^, though several are actively in development by us and others^56^. Therefore, we utilized two B6 tumor models with beta-2 microglobulin deletion^57^. Rejection of allogeneic tumor models is due largely to expression of non-syngeneic major histocompatibility receptors (MHC) and we hypothesized that tumor models with a deficit of MHC class I expression would grow in immune competent NOD mice. In addition, MHC class I deficient tumors are commonly seen in patients since loss of MHC class I expression is a frequent mechanism of immune escape^57–59^. Groups of NOD mice were inoculated with B16.B2M^-/-^ melanoma or MC38.B2M^-/-^ colon carcinoma tumor cells and then treated with ICI or isotype control antibodies. Both tumor models successfully grafted to palpable tumors in NOD mice (**Extended data Fig. 6a**), thus providing two cancer cell lines for testing ICI effects in NOD mice.

Intrathyroidal pan-CD3^+^ IL-17A^+^, and Th17, Tc17, and γδT17 subsets (Fig. 6a), as well as overall CD45^+^ and CD3^+^ immune cells (Fig. 6b), were quantified in groups of tumor-bearing ICI-treated mice after 4 weeks using flow cytometry. In mice with B16.B2M^-/-^ or MC38.B2M^-/-^ tumors, intrathyroidal CD3^+^ IL-17A^+^ cells were increased with Dual ICI-treatment compared to isotype control (p<0.05 and p<0.005, respectively, Fig. 6a). In mice with B16.B2M^-/-^ tumors, Th17 and Tc17 cells were increased (p<0.05 for both), with a trend for increased γδT17 cells (p=0.05). Th17 cells were also increased with Dual ICI-treated mice bearing MC38.B2M^-/-^ tumors (p<0.05). In summary, in tumor-bearing Dual ICI-treated NOD mice, we again see accumulation of IL-17A^+^ T cells in thyroid immune infiltrates, with a significant increase seen primarily in the Th17 subset.

**Figure 6.**
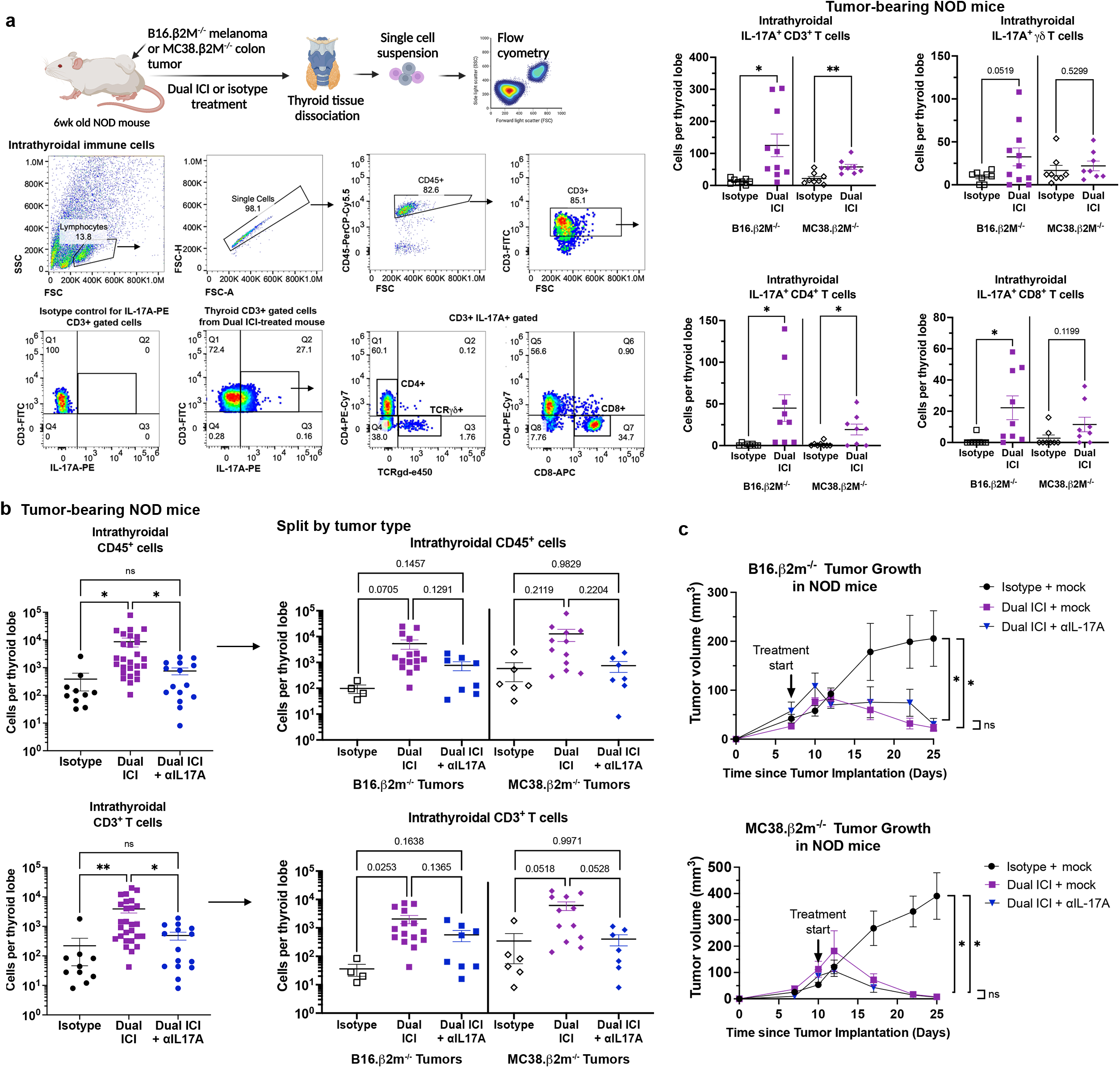
Contribution of IL-17A^+^ T cells in ICI-associated thyroid autoimmune infiltrates in tumor-bearing NOD mice. **a,** Schematic of ICI treatment and thyroid tissue processing in tumor-bearing NOD mice (*top left*). Representative dot plots and gating strategy for thyroid infiltrating immune cells (*bottom left*) Quantification of intrathyroidal IL-17A^+^ CD3+ T cells and IL-17A^+^ T cell subsets in tumor-bearing NOD mice by flow cytometry (*right*). Data shown as cells per thyroid lobe. Each dot represents one animal. **b**, Comparison of accumulation of intrathyroidal CD45^+^ and CD3+ immune cells in isotype, Dual ICI, or Dual ICI with neutralizing IL-17A (αIL17A) antibody therapy, by flow cytometry. Data shown as cells per thyroid lobe. Each dot represents one animal. Summative data shown (*left*) and stratified by tumor model (*right*). **c,** Growth of MC38.β2M^-/-^ and B16.β2M^-/-^ tumors in NOD mice treated with isotype for ICI (isotype) and isotype for αIL17A (mock) (*n*=11-13), Dual ICI with mock (*n*=7), or Dual ICI with αIL17A (*n*=8). Data are mean ±SEM shown. *p<0.05, **p<0.01, ***p<0.001, ****p<0.0001. Two-tailed, unpaired t test with Welch correction, assuming unequal s.d. and Holm-Sidak method correction for multiple comparisons (**a**); Brown-Forsythe ANOVA, assuming unequal s.d., followed by Dunnett’s multiple comparisons test (**b**; comparison of day 25 tumor volume in **c**).

Strategies for blocking IL-17A have been developed for multiple autoimmune indications and could be readily applied to ICI-associated IrAEs. We therefore tested whether IL-17A neutralizing antibody therapy could reduce thyroid autoimmune infiltrates during Dual ICI treatment in tumor-bearing mice. Groups of NOD mice inoculated with B16.B2M^-/-^ melanoma or MC38.B2M^-/-^ colon tumors were treated with Dual ICI or isotype, and IL-17A neutralizing antibody (αIL17A) or isotype (mock), and thyroid infiltrating immune cells were assessed by flow cytometry. As expected, this IL-17A neutralizing antibody reduced the frequency and severity of thyroid autoimmune infiltrates in ICI-treated mice at 4 weeks (Fig. 6b; data shown pooled for statistical analysis and stratified by tumor type). Specifically, thyroid infiltrating CD45^+^ immune cells were reduced from mean 8,636 ± SEM 3,060 cells/lobe in Dual ICI-treated mice, to 761±213 in Dual ICI with αIL17A, and compared to 387±244 in isotype control (ANOVA p=0.0041; p values *vs.* isotype were Dual ICI 0.0362 and Dual + αIL17A 0.5864; p value Dual ICI vs Dual + αIL17A 0.0475). Thyroid infiltrating CD3^+^ immune cells were reduced from mean 3,888±1,054 cells/lobe in Dual ICI-treated mice, to 491±147 in Dual ICI with αIL17A, and compared to 221±175 in isotype control (ANOVA p=0.0004; p values *vs.* isotype were Dual ICI 0.006 and Dual + αIL17A 0.579; p value Dual ICI vs Dual + αIL17A 0.0106). These data support a pathogenic role for IL-17A^+^ T cells in the development of Dual ICI-associated thyroid autoimmune infiltrates and potential benefit for IL-17A blocking therapy in reducing thyroid immune infiltrates.

### Role of IL-17A^+^ T cells in ICI-associated autoimmune infiltrates in other tissues

Similar to cancer patients who often develop IrAEs across multiple tissues^5^, ICI-treated NOD mice develop autoimmune infiltrates in multiple organs (Fig. 1b). Given the role of IL-17A in ICI-thyroiditis and multiple spontaneous autoimmune diseases, we hypothesized that IL-17A may contribute to the development of ICI-associated immune infiltrates in other tissues. IL-17A has previously been implicated in the development of inflammatory bowel disease, a spontaneous autoimmune disease affecting the gastrointestinal system^47^. Importantly, colitis is the second most common IrAE seen in patients, manifesting with diarrhea, weight loss, bloody stool, and even gut perforation^3, 7, 19^. As such, it represents a major clinical challenge to ICI use. In our ICI-treated mice, gut autoimmunity may manifest as weight loss, bloody stools, and gut immune cell infiltration. We assessed the presence of IL-17A^+^ T cells in the gut of ICI-treated NOD mice using flow cytometry (**Extended Data Fig. 6b,** gating strategy same as for thyroid specimens). IL-17A^+^ CD3^+^ T cells were present in the gut of isotype and Dual ICI-treated mice in tumor bearing mice, comprising approximately 15-20% of colon CD45^+^ cells (**Extended Data Fig. 6b**). IL-17A^+^ T cells were comprised primarily of CD4^+^ Th17 cells, with rarer Tc17 and γδT17 cells. Interestingly, while there was not a significant difference in the frequency of IL-17A^+^ T cells in the colon between Dual ICI and isotype treated, treatment with an IL-17A neutralizing antibody significantly reduced the frequency and extent of colon autoimmune infiltrates (**Extended Data Fig. 6c**). This may reflect changes in activation, rather than accumulation or proliferation, of IL-17A^+^ T cells in the colon with ICI treatment.

The NOD mouse develops spontaneous autoimmune diabetes mellitus, and ICI treatment accelerated insulitis and the onset of diabetes. Rare IL-17A^+^ CD3^+^ cells were found in the pancreas of both isotype and ICI-treated mice with no significant difference seen between conditions (**Extended Data Fig. 6d**). Neutralizing IL-17A therapy did not delay the onset of autoimmune diabetes (**Extended Data Fig. 6e**) in ICI-treated mice. In summary, these data suggest that IL-17A contributes to the development of ICI-associated autoimmunity in a tissue-specific manner.

### IL-17A neutralizing antibody during ICI treatment does not reduce anti-tumor effects

Our data suggested that a neutralizing IL-17A antibody may reduce ICI-associated thyroid autoimmune infiltrates. In developing therapies to prevent or reduce IrAE in patients, it will be critical that any approach preserve or enhance the anti-tumor immune effects of ICI. Whether IL-17A inhibition may have deleterious effects on the efficacy of ICI treatment in anti-tumor immunity is unclear, since IL-17A has been associated with pro-tumorigenic effects in some cancers^54, 60, 61^ and anti-tumorigenic effects in others^54, 60, 61^. To test this, we determined the effect of IL-17A blockade during ICI treatment on the growth of tumors in NOD mice. Dual ICI treatment of B16.β2M^-/-^ melanoma and MC38.β2M^-/-^ colon carcinoma in NOD mice significantly reduced tumor growth (Fig. 6c). Importantly, the addition of αIL17A did not reduce the anti-tumor effect of ICI treatment (Fig. 6c; ANOVA for day 25 tumor volume in B16.β2M^-/-^ p=0.0029; p values for comparison to isotype were Dual ICI 0.0226, Dual ICI + αIL17A 0.0286. No significant difference between Dual ICI *vs.* Dual ICI + αIL17A, p=0.9403; ANOVA comparing day 25 tumor volume in MC38.β2M^-/-^ p=0.0004; p values for comparison to isotype were Dual ICI 0.0043, Dual ICI + αIL17A 0.0042. No significant difference between Dual ICI *vs.* Dual ICI + αIL17A, p=0.94). Neutralizing IL-17A antibody therapy combined with Dual ICI treatment of syngeneic MC38 colon tumors in B6 mice also showed no loss of anti-tumor efficacy compared to Dual ICI with a mock isotype control (**Extended Data Fig. 6f**). In summary, blockade of IL-17A function by a neutralizing antibody during ICI treatment significantly reduced ICI-associated autoimmune infiltrates while preserving the anti-tumor efficacy of anti-PD-1 + anti-CTLA-4 therapy.

## DISCUSSION

Immune checkpoint inhibitors (anti-PD-1/L1 and anti-CTLA-4) have significantly advanced the treatment of cancer since their first approval in 2011. However, the benefits and use of ICI have been limited by the frequent development of unwanted IrAEs that contribute to patient morbidity and may lead to interruption of cancer treatment. Despite significant efforts to date, the cause of IrAEs remained poorly understood. Data from thyroid specimens from patients highlighted a role for diverse T cell populations in ICI-thyroiditis, including a rare population of *RORC^+^IL23R^+^* T cells related to Type 3 immunity and classically associated with Hashimoto’s thyroiditis^24–26, 49^. Using a novel mouse model in which checkpoint inhibitor therapy leads to multi-organ autoimmune infiltrates, we identify Type 3 immune cells including γδT17 and Th17 cells as critical contributors to IrAE development in this model. Indeed, antibody-based inhibition of IL17A protected mice from ICI-induced autoimmune infiltrates. Finally, IL-17A neutralization during ICI treatment did not reduce ICI anti-tumor efficacy. Thus, targeting Type 3 immune cells in ICI-treated patients may reduce IrAE without impairing the anti-tumor efficacy of ICI.

In our model, ICI-treated NOD mice developed thyroiditis comprised of a T cell predominant but diverse immune infiltrate. Angell *et al.* first reported on the thyroid immune cell infiltrate in a patient with ICI-thyroiditis, noting lymphocytic cells and histiocytes^14^. In a subsequent study, Kotwal *et al.*^62^ evaluated immune cells in thyroid FNA specimens from eight ICI-thyroiditis patients and noted a predominant CD3^+^ T cell population, as well as myeloid cells. This study demonstrated an increase in PD-1^+^ T cells in thyroid immune infiltrates compared to healthy controls that is similar to our findings from scRNAseq data showing increased checkpoint protein (e.g. *Pdcd1, Tigit, Lag3*) gene expression by thyroid-infiltrating T cells in specimens from ICI-thyroiditis patients and ICI-treated mice. Such intrathyroidal T cell populations may be direct targets of ICI. Interestingly, Kotwal et al. also noted a significantly increased CD4^-^ CD8^-^ CD3^+^ T cell population in ICI-thyroiditis patients compared to healthy controls (1.9% *vs.* 0.7% of CD45^+^ cells), though these were not characterized as γδ T cells. Our data similarly support the accumulation of intrathyroidal γδT cells in ICI-thyroiditis. Finally, a recent report by Yasuda *et al.*^33^ evaluating anti-PD-1 therapy in mice pre-immunized with human thyroglobulin, showed a key role for CD4^+^ T cells in the development of ICI-associated thyroiditis. While our results are consistent with this, our studies further demonstrate the importance of Th17 subsets within CD4^+^ T cells as well as non-CD4^+^ γδT17 subsets.

While Th17 cells are well described in spontaneous autoimmunity, including Hashimoto’s thyroiditis^24–26^, they have not previously been reported in IrAE. Our data highlight a shared role for Th17 cells in these two forms of thyroiditis. γδT17 cells, in contrast, have not previously been described in studies of spontaneous thyroiditis or in IrAE^63^. γδT17 cells are unique in their early activation and robust cytokine production compared to αβ TCR T cells^52, 64, 65^. In addition, γδT17 cells have been reported to expand Th17 cells, and thus early γδT17 activation may contribute to activation and recruitment of Th17, CD8^+^, and other T cell populations. Future studies can explore the role of γδT17 cells in spontaneous and ICI-induced thyroiditis.

The IrAE predisposition of the inbred NOD strain may mimic humans with genetic predisposition to IrAE development. For instance, HLADR15 haplotype has been associated with pituitary IrAE^66^ while the HLADR4 haplotypes has been associated with autoimmune diabetes IrAE in humans^21^. Autoimmune predisposition has been mapped to NOD polymorphic regions which include MHC. This model using NOD mice overcomes some critical limitations of previous models that required deletion of regulatory populations, transfection of human checkpoint proteins, or the requirement for chemical stimulants or xeno-antigen immunization for autoimmunity^23, 30–33^. For instance, NOD.H2h4 mice, which have been used previously to study thyroiditis^24, 67, 68^ and anti-CTLA-4 induced thyroiditis^32^, require excess dietary NaI to precipitate thyroiditis via follicular cell injury and TNFα stimulation, have high background rates of thyroiditis (100% after 6 weeks of NaI supplementation^67^), and require the presence of a non-syngeneic MHC class II molecule^42, 67^. In contrast, our model in NOD mice provides an immune competent system with low rates of spontaneous autoimmunity at advanced age, more similar to the background human population. The background incidence of spontaneous thyroiditis of approximately 14% at 1 year in NOD mice^38^ compares to a prevalence of near 10% for underlying thyroid autoimmunity in the general population^69–71^. As seen in humans, mice with a genetic predisposition (*i.e*. NOD *vs*. B6) developed more IrAE across multiple tissues following ICI treatment and with combination *vs.* single agent ICI therapy^21, 35^.

While our study focused on thyroid IrAE, a common autoimmune side effect encountered during ICI treatment that results in permanent organ dysfunction, these mechanisms may be shared among IrAE in other organs with more grave clinical consequences. Mice in our model developed multi-organ autoimmunity, including colitis, pneumonitis, hepatitis, and nephritis, and therefore can be used to study IrAE in these organs. Neutralizing IL-17A antibody therapy decreased overall autoimmune infiltrates, including in non-thyroid tissues, and therefore may be of utility in reducing IrAE during ICI more broadly. ICI-associated colitis has been associated with increased circulating IL-17A in patients treated with anti-CTLA-4 ^72, 73^ and IL-17 has previously been shown to contribute to spontaneous gut autoimmune diseases, including inflammatory bowel disease^47^. IL-17A^+^ T cells were prevalent in the colon of NOD mice and colon autoimmune infiltrates were reduced with a neutralizing IL-17A antibody. Future studies into the role of the IL-17A axis in ICI-associated IrAE in the colon and other organs are warranted and are facilitated by this new mouse model.

IrAEs remain a significant barrier to the use of ICI but therapeutic strategies to prevent IrAEs have not been widely implemented. A primary consideration in the development of strategies to reduce IrAEs must be the preservation of anti-cancer effects of ICI. As shown here, our model can facilitate these studies using the B16 melanoma and MC38 colon B6 tumor models with genetic deletion of β2M. Several syngeneic NOD tumor models are also in active development by us and others. Early efforts using glucocorticoids for IrAE treatment inconsistently reversed autoimmunity or compromised anti-cancer effects^5, 20^. In addition, previous studies have shown reduction of thyroid IrAE by elimination of CD4^+^ T cells^33^, but this is not readily translatable to cancer patients given the importance of CD4^+^ T cells in cell-mediated immunity^40, 74^. Similarly, while Tbet1^+^ IFNγ-producing T cells are seen in thyroid immune infiltrates and almost certainly contribute to autoimmunity^75^, these cells are known to be critical to anti-cancer effects of ICI^76–78^ and therefore cannot feasibly be inhibited for IrAE prevention. Here we provide evidence that IL-17A blockade may be useful for reducing IrAE without hindering anti-tumor effects. Therapies targeting the IL-17A pathway are already FDA approved for use in patients. Based upon these results, we propose that a clinical trial evaluating inhibition of the IL-17A axis for prevention of IrAEs in cancer patients receiving ICI treatment should be pursued.

## METHODS

### Mice

NOD/ShiLtJ (NOD) and C57/B6 (B6) mice were obtained from the Jackson Laboratory. Male and female mice were used in equal proportions unless otherwise specified. Mice were used at 4-6-week of age unless otherwise noted. Mice were housed in a specific pathogen-free barrier facility at the University of California Los Angeles. Diabetes, determined by presence of glucose in urine (Diastix, Bayer), was assessed at least once per week and diabetic mice were treated daily with intraperitoneal insulin as described previously^39^ until used in experiments or euthanized. All experiments were conducted under IACUC-approved protocols and complied with the Animal Welfare Act and the National Institutes of Health guidelines for the ethical care and use of animals in biomedical research.

### Cell lines

Cell lines used in these studies included MC38, a murine colon tumor model; MC38 with genetic deletion of beta 2 microglobulin (MC38.β2M^-/-^); murine melanoma tumor model B16 with β2M deletion (B16.β2M^-/-^)^57^; and CHO-mTPO, Chinese hamster ovary cells stably transfected for surface membrane expression of mouse thyroid peroxidase (TPO) protein expression^68^. Tumor cell lines were obtained from the American Type Culture Collection [(ATCC), MC38] or were gifted to Dr. Lechner and Dr. Su (CHO-mTPO cells by Drs. Basil Rapoport and Sandra McLachlan; MC38.β2M^-/-^ and B16.β2M^-/-^ cells by Dr. Antoni Ribas). Tumor cell line authenticity was performed by surface marker analysis performed at ATCC or in our laboratory. MC38, MC38.β2M^-/-^ and B16.β2M^-/-^ cells were grown in complete medium [RPMI-1640 supplemented with 10% fetal bovine serum (FBS), 2mM L-glutamine, 1mM HEPES, non-essential amino acids, and antibiotics (100 U/mL Penicillin and 100 μ Streptomycin)], at 37°C in humidified, 5% CO_2_ incubators. CHO-mTPO cells were cultured in F12 media supplemented with 10% FBS, 2mM L-glutamine, 1mM HEPES, non-essential amino acids, and antibiotics (100 U/mL Penicillin and 100 μg/mL Streptomycin). Cell lines were monitored regularly for phenotype. Early passage cells were used for experiments (P1 or P2).

### Reagents and media

Immune checkpoint inhibitor antibodies used were anti-mouse PD1 (clone RPM1-14), CTLA-4 (clone 9D9), and isotype controls [clone 2A3 and MPC-11, respectively] (BioXcell). For inhibitor experiments, a neutralizing antibody against mouse interleukin-17A (clone 17F3) or isotype control (clone MOPC-11), from BioXCell. Antibodies were diluted in sterile PBS for use. For *in vitro* experiments, primary immune cells were cultured in RPMI complete media [supplemented with 10% fetal bovine serum (FBS), 2mM L-glutamine, 1mM HEPES, non-essential amino acids, and antibiotics (penicillin and streptomycin)], with 50uM beta-mercaptoethanol.

### Immune checkpoint inhibitor treatment of mice

Groups of 4-6 week old NOD or B6 mice were used for ICI inhibitor experiments. Mice were randomized to twice weekly treatment with anti-mouse CTLA-4 (clone 9D9), or PD-1 (RPM1-14), both anti-CTLA-4 and anti-PD-1, or an isotype control (2A3, MPC-11), at 10 mg/kg/dose intraperitoneally *(i.p.)* for four or eight weeks. During treatment mice were monitored daily for activity (including signs of neuropathy) and appearance, and twice weekly for weight and glucosuria. Mice developing glucosuria were treated with 10 units of subcutaneous NPH insulin daily. Thyroid dysfunction was assessed by measurement of free thyroxine (FT4) in sera by ELISA (LSBio). After four or eight weeks of ICI treatment (as indicated), mice were euthanized, blood collected by retro-orbital bleed, then perfused with 10mL of phosphate buffered saline (PBS). Tissues were collected for histologic analysis into neutral buffered formalin, including thyroid, lacrimal and salivary glands, lung, liver, kidney, heart, colon, eye, gonad, and pancreas. Serum was collected from blood (centrifugation 1000x*g* for 30 min). Predetermined endpoints for euthanasia before four weeks included >20% weight loss and glucosuria not resolved by insulin therapy, as per IACUC protocols.

### Histology

Harvested organs were fixed in 10% buffered formalin for at least 96 hours and then stored in 70% ethanol. Organs were embedded in paraffin, sectioned (4 μm), and stained with hematoxylin and eosin (H&E) by the UCLA Translational Pathology Core Laboratory. Immune infiltration was assessed on H&E sections using an adapted immune infiltrate scoring system reported previously^39^. The presence of interstitial inflammation, lymphocytic aggregates, perivascular inflammation, follicular disruption, and Hurthle cell change was assessed as either present or absent on H&E tissue sections by a clinical pathologist (E.D.R.). In addition, each tissue was given an aggregate immune infiltrate score of 0 (no immune infiltrate), 1 (1-2 focal areas of immune infiltrate and/or sparse interstitial inflammation), 2 [>2 focal areas of immune infiltration, and/or presence of both focal glomerulonephritis and perivascular immune infiltration (kidney), and/or diffuse immune infiltrate affecting >25-49% of the tissue area] or 3 (diffuse immune infiltrate affecting >50% of the tissue area). For pancreas tissue, immune infiltrate score was modified as 0 for no immune infiltrate, 1 for <50% of islets affected by immune infiltration, or 2 for >50-100% of islets affected by immune infiltration. The enumeration and typing of inflammatory infiltrate were assessed through immunohistochemical stains (see below). Immune scoring was done by two blinded individuals evaluating at least 10 high powered fields per section for pancreas, lung, liver, heart, thyroid, kidney, salivary, gut (colon and stomach), and gonad tissues; for small tissues (lacrimal glands and eye) 5 high powered fields were evaluated. Images were acquired on an Olympus BX50 microscope using Olympus CellScans Standard software. Images were brightened uniformly for publication in Photoshop.

### Immunohistochemistry

For immunohistochemistry (IHC), 4mm FFPE tissue sections were deparaffinized, rehydrated, and subjected to heat-induced antigen retrieval (0.01 mol/L citrate, pH 6.0) followed by treatment with 3% H_2_O_2_ for 10 minutes to block endogenous peroxidase activity. Sections were incubated overnight at 4°C with primary antibodies against mouse IL17A (Abcam, ab91649, 1:50 dil), CD3 (DAKO, clone A0542, 1:100 dil), F4/80 (BioRad, clone MAC497G, 1:200 dil), B220 (BD, clone RA3-6B2, 1:50 dil). Sections were then stained with appropriate secondary antibodies and antigen detection done with 3,30-diaminobenzidine. Sections were counterstained with hematoxylin, dehydrated, and mounted. Appropriate positive and negative controls were used for all stains. Brightness of all images was increased by 50 in Adobe Photoshop. Positively stained leukocytes for CD3, B220, F4/80, or IL-17A were counted in 10 representative high-power fields (hpf, total 400x magnification) for each tissue section. Two independent observers scored each section and the results were pooled with rare disagreements resolved by a third evaluator.

### ELISA for thyroglobulin antibody

Mouse anti-thyroglobulin antibodies were measured via ELISA as previously reported^42^. Briefly, 96-well plates were coated with 50uL of 1.5mg/mL purified mouse thyroglobulin (mTg, generously provided by the Rapoport lab) in Coating Buffer (30mM Na2CO3, 70mM NaHCO3, 6mM NaN3, pH 9.3) covered and incubated at 4°C overnight. The following day, plates were washed twice with 100uL per well Tris/NaCl (200mM Tris base, 188mM NaCl, pH 7.4), once with Tris/NaCl Tween (Tris/NaCl, 500uL/L Tween 20), and once more with Tris/NaCl. Plates were blocked with 100uL/well Blocking Buffer (5% w/v BSA, 150mM NaCl) for twenty minutes at room temperature (RT) then washed again as above. Sera samples were diluted 1:100 in PBS and added to wells in triplicate. Plates were covered and allowed to incubate RT 1.5 hours. Plates were washed four times with 100uL/well of Tris/NaCl Tween. Anti-mouse IgG1-HRP was diluted 1:2000 in Secondary Antibody Buffer (1% w/v BSA, 150mM NaCl) and was added to plate at 100uL/well. Plate was covered and allowed to incubate RT one hour, then washed four times with 100uL/well Tris/NaCl Tween. ELISA was read by adding 100uL/well TMB substrate (Thermo Scientific, N301), allowing 10 minutes to develop in dark, and read on a spectrophotometer at OD 605nM.

### Thyroid peroxidase antibody

Mouse thyroid peroxidase antibody (TPO) antibody levels were detected using mouse sera binding on CHO-mTPO cells, followed by flow cytometry as previously described^42^. Briefly, experimental mouse sera were incubated with a single cell suspension of mouse CHO-mTPO cells at a 1:50 dilution in Buffer A (2% FBS in PBS with 10mM HEPES) for 45 minutes at 4°C, then washed twice with Buffer A. For detection of immunoglobulin binding, samples were then incubated with goat anti-mouse IgG FITC secondary antibody at 10ug/mL for 30 minutes at 4°C and then washed twice with Buffer A. Finally, mean fluorescence intensity of FITC was measured for live, single CHO-cells using an Attune NxT 6 cytometer (ThermoFisher).

### Flow Cytometry

Immediately after euthanasia and perfusion with sterile PBS, fresh tissues were dissociated for analysis of immune infiltrates by flow cytometry. Thyroid glands were dissected away from surrounding trachea and lymphoid tissue, digested in collagenase type IV (1mg/mL in 2% FBS in PBS) at 37°C for 20 minutes, then mechanically dissociated by passage through a 40um filter. Spleen cells were isolated by mechanical dissociation and passage through a 40um filter. To assess intracellular cytokines, cells were incubated in complete RPMI media with 50uM 2ME for four hours with ionomycin (1ug/mL) and PMA (50ng/mL) in the presence of Brefeldin A prior to staining. For staining, single cell suspensions were resuspended in FACS buffer (0.5mM EDTA, 2% FBS in PBS) at 10^6^cells/mL and stained with fluorescence conjugated antibodies. For intracellular staining, after surface staining, cells were fixed and permeabilized using cytoplasmic fixation and permeabilization kit (BD, for cytokine IL-17A) or FoxP3 transcription factor kit (eBioscience, for RORγt), per protocol instructions, with 30 min fixation at 4°C. Viability dye DAPI was added prior to analysis where indicated. Cells were then washed twice in FACS buffer and analyzed by flow cytometry on an Attune NxT 6 cytometer (ThermoFisher). Antibodies used are shown in **Supplemental Methods**. Cell counts are shown as relative frequency of live, gated single cells unless otherwise noted. For determination of infiltrating cells per thyroid lobe, absolute cell counts per thyroid lobe were determined by a calculation of cell count x fraction of thyroid analyzed to estimate a total cell count per thyroid lobe. Whole perfused thyroid specimens were dissociated to single cell suspensions and the entire specimen stained for flow cytometry and a fixed volume of the specimen was evaluated by flow cytometry to allow back calculation for estimate of absolute cell count (*e.g.* 100ul of 200ul total sample volume run yields a 2x multiplier for cell count). Typical thyroid lobe weight ranged from 0.2-0.4mg.

Immune cells were isolated from gut tissue mice to assess colitis as previously described^79^. Briefly, colons were dissected from each mouse after euthanasia and perfusion as above, and cut into 1 cm pieces. Tissue pieces were first washed with 10% FBS + 2mM EDTA in PBS, underwent further mechanical dissociation with a razor blade and then digested in 1 mg/ml collagenase type IV at 37°C for 20 minutes. After digestion, cells were washed and rested in complete RPMI media to quench remaining collagenase present. Lymphocytes were isolated using Percoll gradient centrifugation. Lymphocytes were collected, washed with PBS, and used for flow cytometry analysis as above.

### Neutralizing antibody studies in ICI-treated mice

Groups of 4-6 wk old NOD mice were randomized to treatment with anti-mouse PD-1 (clone RPM1-14) plus anti-mouse CTLA-4 (9D9) antibodies or isotype controls (2A3 plus MPC-11) at 10mg/kg/dose twice weekly by *i.p.* injection. Additionally, 10 days after start of ICI or isotype therapy, mice were further randomized to receive neutralizing antibody to IL-17A (clone 17F3) or isotype (clone MOPC-11).

### Tumor model studies

Groups of 6-week old C57/B6 mice were inoculated *s.c.* with 3×10^5^ MC38 tumor cells in the flank, as described previously^80^. Similarly, 6-week old NOD mice were inoculated *s.c.* with 6×10^5^ MC38.β2M^-/-^ or B16.β2M^-/-^ tumor cells in the flank. Mice were randomized into groups for treatment when tumor volumes reached 40–80mm^3^. Groups of mice received anti-mouse PD-1 (clone RPM1-14), anti-mouse CTLA-4 (9D9), both, or isotype controls (2A3 plus MPC-11) at 10mg/kg/dose twice weekly, alone or in combination with a neutralizing antibody to mouse IL-17A (clone 173F, 0.5mg/dose, three times weekly) or isotype control (clone MOPC-11). Reagents were given by *i.p.* injection on the ipsilateral side to the tumor. Mouse tumor volumes were measured every 2 days by caliper and mice were sacrificed when tumor volumes exceeded 2cm in diameter or when animal morbidity mandated sacrifice under institutional vivarium protocols.

### Quantitative reverse transcriptase PCR (qRT-PCR)

For RNA extraction, human NThy-Ori-3.1 cells were washed twice with PBS, then lysed directly in tissue culture wells. For mice experiments, resected mouse thyroid lobes were snap frozen in liquid nitrogen and stored at −80°C. Multiple thyroid lobes were homogenized in RNA Lysis Buffer (Zymo Research) using the Fisherbrand 150 Handheld Homogenizer (15-340-167). RNA extractions were performed using Quick-RNA Microprep kit (Zymo Research, R1051) per manufacturer’s instructions. RNA yield was quantified by nanodrop. RNA was converted into cDNA using High-Capacity cDNA Reverse Transcription Kit with RNAse Inhibitor and MultiScribe Reverse Transcriptase (Applied Biosystems, 4374966) following manufacturer’s instructions. About 750 ng/uL of RNA was used for cDNA synthesis per reaction. Quantitative PCR was performed using Taqman Fast Advanced PCR Master Mix (Applied Biosystems, 4444557) and Taqman probes for genes of interest (**Supplemental Methods**), with three technical replicates per gene, per condition. PCR cycles were run using the QuantStudio 6 Pro PCR machine (Applied Biosystems) with the standard cycle parameters. Design and Analysis QuantStudio 6/7 Pro Systems Software (Thermo Fisher Scientific, Version 2.5.0) was used to identify amplification of genes and calculate fold change from Cq values. In this context, fold change was the expression ratio of the gene of interest to the housekeeping gene, GAPDH. For further analysis of NThy-Ori-3.1 antigen presentation and cytokine production, human 293T cells were also used for qRT-PCR to compare upregulated gene expression.

### Patients

Patients were prospectively enrolled from two academic medical centers (UCLA Health, USC Keck Medical Center) under IRB approved protocols (19-001708, HS-19-00715). Thyroid FNA specimens were collected from adult (age >18 years) patients with 1) ICI-treated cancer patients with new onset thyroid IrAE, or 2) Hashimoto’s thyroiditis (HT). Exclusion criteria included pregnancy, history of thyroid surgery, radioactive iodine therapy, or thyroid cancer, immune modifying conditions not including solid malignancy (e*.g.* bone marrow transplantation, leukemia or lymphoma, known genetic or acquired immunodeficiency, or immune modifying medications at the time of specimen collection, excluding physiologic steroids). ICI-thyroiditis (Thyroid IrAE) was defined as new onset thyroid dysfunction (within 2 months of specimen collection) while on anti-PD-1/L1 and/or anti-CTLA inhibitor therapy, with an overt hyperthyroid phase (low TSH and elevated FT4) followed by hypothyroidism (elevated TSH and low FT4) requiring thyroid hormone replacement. ICI-treated subjects must have received an FDA-approved ICI within the past 1 month. HT patients had hypothyroidism (elevated TSH and low FT4, and/or requirement for thyroid hormone replacement) and evidence of thyroid autoimmunity (e*.g.* thyroid autoantibody presence, anti-TPO or anti-Tg); imaging findings if available were consistent with HT. Patients may have had a cytologically-proven benign thyroid nodule. Patient demographic, treatment and clinical immune data are summarized in **Extended Data Table 1**. Thyroid FNA specimens were collected into RPMI media under ultrasound guidance using 4 passes with a 25 or 27 gauge needle by M.G.L, T.E.A., or P.F.

### Single cell RNA sequencing

For mouse scRNAseq studies, CD45^+^ infiltrating cells were isolated from fresh thyroid tissue. To reduce contamination from circulating immune cells in the blood, animals were perfused with normal saline prior to tissue collection. Thyroid tissue was enzymatically and mechanically dissociated, then single cell suspensions stained with fluorescence-conjugated antibodies to CD45, CD11b, CD3, CD4, CD8, and CD19 and viability dye DAPI. Specimens with and potential cervical thymus contamination were excluded before sequencing using a threshold of >20% double positive CD4^+^ CD8^+^ cells within CD3^+^ T cells^81^. Samples for each condition were then pooled to obtain sufficient cells for sequencing and analysis. Cell preparation, library preparation, and sequencing were carried out according to Chromium product-based manufacturer protocols (10X Genomics). Sequencing was carried out on a Novaseq6000 S2 2×50bp flow cell (Illumina) utilizing the Chromium single-cell 5′ gene expression library preparation (10X Genomics), per manufacturer’s protocol at the UCLA Technology Center for Genomics and Bioinformatics (TCGB) Core. Data were demultiplexed and aligned with Cell Ranger version 3.0.0 or higher (10X Genomics)

For human scRNAseq of PBMC and thyroid FNA specimens, sorted, single live CD45^+^ immune cells were similarly collected and sorted, then submitted for 10x sequencing. Library construction and sequencing was done by the UCLA Technology Center for Genomics and Bioinformatics (TCGB) core facility.

### Data processing of scRNA-seq libraries

After sequencing, the scRNA-seq reads were demultiplexed and aligned with Cell Ranger version 3.0.0 or higher (10X Genomics) to the mm10 (mouse) or GRCh38 (human) reference genome and quantified using the 10x Genomics CellRanger count software (10x Genomics). Filtered output matrices that contained only barcodes with unique molecular identifier (UMI) counts that passed the threshold for cell detection were used for downstream analysis.

### Normalization, Principal Component analysis, and UMAP Clustering

Downstream analysis was done with Seurat v3 or higher^82^. Only cells with a mitochondrial gene percentage less than 30% and 200 features were included in downstream analysis. Scores for S and G2/M cell cycle phases were assigned using the Seurat CellCycleScoring function following the standard Seurat pipeline^83^. UMI counts were log normalized, and the top 2000 variable genes were determined using the variance-stabilizing transformation (vst) method. All genes were scaled and centered using the ScaleData function, and principal component analysis (PCA) was run for the data using the predetermined variable genes. To group cells into clusters, a K-nearest neighbors graph function (implemented in the Seurat package), followed by a modularity-optimizing function using the Louvain algorithm was used. For the cluster, 30 PC dimensions were included and the resolution parameter was set to 0.4. Cell-type clusters were visualized using uniform manifold approximation and projection (UMAP) to reduce dimensionality and allow for the cells to be visualized on a 2-D plot.

### Differential Expression and Marker Gene Identification

The Seurat FindMarkers function was used to generate the top upregulated genes for each cluster using a Wilcoxon Rank Sum Test to identify differentially expressed genes across clusters. Marker genes were filtered by a minimum of detectable expression in 25% of the cells in the target group and minimum log2 fold change of 0.25. The markers generated by these functions were compared to markers for known cell types to assign identities to the different clusters. Specifically, variably expressed gene sets for each cluster were queried in curated, publicly available databases for putative cell population identity: Enrichr^29^ (https://maayanlab.cloud/Enrichr) and the Immunological Genome Project database^45^ (http://rstats.immgen.org/MyGeneSet_New/index.html). Differentially expressed genes across conditions were identified using the same function and parameters. Cells from proliferating clusters, stressed/dying clusters, thymic contamination, doublets, and unknown clusters were excluded from this analysis.

### Statistics

Data were analyzed with GraphPad Prism v9. Descriptive statistics shown are mean + SEM for continuous variables. Differences in the mean frequency of immune cells or gene expression for qRT-PCR between two groups were evaluated by two-sided t test with Welch correction for potential differences in variance among groups and correction for multiple comparisons by the Holm-Sidak method. For B6 mice, differences in organ immune infiltrate score, thyroid autoantibodies (TPO MFI and Tg ELISA OD) were compared between groups by two-tailed, unpaired t test with Welch correction, without assumption of equal s.d. Differences in the mean frequency of immune cells, organ immune infiltrate score, thyroid autoantibodies (TPO MFI and Tg ELISA OD) among groups were evaluated by Brown-Forsythe ANOVA with Welch correction for potential differences in variance among groups, followed by multiple comparisons between groups with Dunnett correction for multiple comparisons (two-tailed comparisons). Where appropriate, adjusted p values are shown. Significance was set at an alpha = 0.05. Samples sizes for each group or condition are shown.

### Data availability

Data associated with figures are available from the corresponding author upon reasonable request. The datasets for single cell sequencing generated during and analyzed during the current study are available in the Gene Expression Omnibus (GEO) repository under accession number GSE192581 (https://www.ncbi.nlm.nih.gov/geo/).

## Acknowledgements

This work was supported by funding from the American Thyroid Association (THYROIDGRANT2020-0000000169, M.G.L.), National Institutes of Health (K08 DK129829-01, M.G.L.), Aramont Charitable Foundation (M.G.L.), Parker Institute for Cancer Immunotherapy (M.A.S.), and UCLA core facilities [Broad Stem Cell Research Center FACS core, Translational pathology core laboratory (TPCL), and Technology Center for Genomics and Bioinformatics (TCGB)]. The authors would like to acknowledge Drs. Sandra McLachlan and Basil Rapoport for CHO-mTPO cell line and methods for thyroid autoantibody detection.

## Author contributions

M.G.L., W.H., T.E.A., A.D., and M.A.S. conceived the study and designed the analysis. M.G.L., A.Y.P., T.E.A., M.S.P., N.Y., M.I.C., A.T.H., L.G., M.A, and P.F. collected the data, including mouse experiments with ICI treatment, patient specimens, flow cytometry, qRT-PCR, immunohistochemistry, and tumor model studies. W.H., A.R., A.D., H.C., and M.A. S. contributed data and analysis tools. M.G.L., A.Y.P., M.S.P., N.Y., A.T.H., E.C.M., E.D.R., and H.C. performed the analysis. M.G.L., A.Y.P., W.H., T.E.A., M.A.S., A.T.H., H.C., N.Y., E.C.M., A.D., A.R., and M.A.S. wrote or edited the paper. All authors reviewed the manuscript.

## Competing interests

A.R. has received honoraria from consulting with Amgen, Bristol-Myers Squibb, Chugai, Genentech, Merck, Novartis, Roche, Sanofi and Vedanta, is or has been a member of the scientific advisory board and holds stock in Advaxis, Appia, Apricity, Arcus, Compugen, CytomX, Highlight, ImaginAb, Isoplexis, Kalthera, Kite-Gilead, Merus, PACT Pharma, Pluto, RAPT, Rgenix, Synthekine and Tango, has received research funding from Agilent and from Bristol-Myers Squibb through Stand Up to Cancer (SU2C), and patent royalties from Arsenal Bio. All other authors have declared no conflicts of interest.

## Additional information

Extended data is available for this paper.

Supplementary information is available for this paper.

Correspondence and materials requests should be directed to Dr. Melissa Lechner, Division of Endocrinology, Diabetes, and Metabolism, UCLA, 10833 Le Conte Ave, CHS 57-145, Los Angeles, CA, 90095.; Phone: 310-794-3237.

## Extended Data Tables

Extended Data Table 1. Demographic and clinical data for human subjects with thyroid disease.

Extended Data Table 2. Variable expression gene lists for scRNAseq data.

## Extended Data Figure Legends

**Extended Data Fig. 1. Supplemental data for flow cytometry and scRNAseq of human thyroid fine needle aspiration (FNA) specimens from Hashimoto’s thyroiditis (HT) and immune checkpoint inhibitor (ICI)-thyroiditis patients. a,** Gating strategy and representative dot plots for immune cells. b, The expression levels of various marker genes of each cluster in human thyroid specimen UMAP clusters for relevant immune populations.

**Extended Data Fig. 2. Flow cytometry analysis of peripheral immune changes in ICI-treated NOD mice**. **a,** Relative frequency of putative T (CD3^+^ NKp46^-^), natural killer (NK, NKp46^+^ CD3^-^), NKT (CD3^+^ NKp46^+^), B (B220^+^ CD3^-^ NKp46^-^ CD11b^-^), myeloid (CD11b^+^) and conventional dendritic cell (DC, CD11c^+^ CD11b^low^) as a percent of splenocytes in isotype (*n*=7) *vs.* anti-PD-1 + anti-CTLA-4 (Dual ICI, *n*=8) mice after 4 weeks of treatment. P values are as follows: T cell 0.012, NK cell 0.004, NKT cell 0.002, B cell 0.12, CD11b^+^ myeloid cell 0.3, and DC 0.16. **b,** Percent of CD4^+^, CD8^+^ and γδTCR^+^ (γδ) T cells in spleens of Dual ICI-treated (*n*=8) or isotype (*n*=8) mice expressing activation markers (CD44^+^ CD62L^-^), by flow cytometry. P values are as follows: CD4^+^ 0.002, CD8^+^ 0.0003, γδ 0.29. d, Relative frequency of regulatory T cells within CD3+ cells in spleen of ICI-treated (*n*=8) or isotype (*n*=7) mice, p=0.0001. Data shown are mean ± SEM. *p<0.05, **p<0.01, ***p<0.001. Differences in immune populations in spleen were compared by two-tailed, unpaired t test with Welch correction, assuming unequal s.d., and Holm-Sidak method correction for multiple comparisons.

**Extended Data Fig. 3. ICI therapy in B6 mice induces minimal tissue autoimmunity.**

**a**, Schematic of ICI drug treatment. **b**, Comparison of autoimmune organ infiltration after 4 weeks of ICI *vs.* isotype treatment in C57/B6 (B6) mice (*n*=8 isotype, *n*=8 Dual ICI); Pie charts (*right*) show tissue infiltrate for each mouse; black = immune infiltrate, white = no infiltrate. **c,** Flow cytometry analysis of thyroid-infiltrating immune cells in anti-PD-1 + anti-CTLA-4 (Dual ICI) treated B6 mice. Mice were euthanized after 4 weeks of ICI or isotype treatment, perfused with saline, and then fresh thyroid tissues dissociated into single cell suspensions. Cells were stained with fluorescent antibodies and analyzed by flow cytometry, with estimation of cells/thyroid lobe for each animal. Each point represents an individual mouse (*n*=8 isotype and *n*=8 Dual ICI). **d**, Quantification of anti-thyroid autoantibodies in B6 mice after 4 weeks of isotype (*n*=8) or Dual ICI (*n*=8) treatment. Data are mean±SEM. Groups were compared by two-tailed, unpaired t test with Welch correction, assuming unequal s.d., and Holm-Sidak method correction for multiple comparisons (**b, c, d**). *p<0.05.

**Extended Data Fig. 4. Supplemental data for scRNAseq of thyroid immune infiltrates from isotype or Dual ICI-treated NOD mice**. **a**, The expression levels of various marker genes of each cluster in mouse thyroid tissue UMAP clusters for relevant immune populations. **b,** Expression of immune genes associated with Type 3 immune responses across cell clusters.

**Extended Data Fig. 5. Gating strategy and representative dot plots for primary mouse immune cells. a,** Gating strategy and representative dot plots for CD45^+^, CD3^+^, Nkp46^+^, B220^+^, CD11c^+^ CD11b^+^, and CD11c^+^ CD11b^low^ cells. **b,** Gating strategy and representative dot plots for CD4^+^, CD8^+^, and TCRγδ^+^ T cell subsets. **c,** Representative dot plots for IL-17A^+^ staining of CD45^+^ immune cells showing CD3^+^ and CD3^-^ populations, and variability in the amount of intrathyroidal immune cells across ICI-treated specimens (*center* and *right* panels).

**Extended Data Fig. 6. Tumor growth and ICI-associated IL-17A^+^ T cells in non-thyroid tissues in NOD mice. a,** Representative growth curves for tumors in NOD mice inoculated with or B16.β2M^-/-^ melanoma or MC38.β2M^-/-^ colon carcinoma tumor cells. **b,** Relative frequency among CD45^+^ immune cells of IL-17A^+^ CD3^+^ T cells and IL-17A^+^ T cell subsets isolated from colon tissue of isotype or Dual ICI-treated mice at 4 weeks by flow cytometry, 3 experiments, each dot represents a single animal. **c**, Comparison of colon immune infiltrate on histology for isotype (*n*=11), Dual ICI-treated (*n*=11), or Dual ICI and neutralizing IL-17A (αIL17A) therapy (*n*=14) NOD mice after 4 weeks, 2 experiments, each dot represents a single animal. **d**, Relative frequency among CD45^+^ immune cells of IL-17A^+^ T cells in the pancreas of isotype *vs.* Dual ICI-treated mice, 2 experiments, each dot represents a single animal. **e,** Kaplan-Meier curve showing incident autoimmune diabetes mellitus in NOD mice among treatment groups as assessed by persistent glucosuria requiring insulin therapy. **f,** Growth of syngeneic MC38 colon tumors in C57/B6 (B6) mice treated with isotype, Dual ICI, or Dual ICI and αIL17A, *n=*6-8 each group. Data are mean ±SEM shown. *p<0.05, **p<0.01, ***p<0.001, ****p<0.0001. Brown-Forsythe ANOVA, assuming unequal s.d., followed by Dunnett’s multiple comparisons test (**b,c**); Two-tailed, unpaired t test with Welch correction, assuming unequal s.d. and Holm-Sidak method correction for multiple comparisons (**d**); Log rank test for trend for median diabetes-free survival (**e**); or ANOVA for tumor volume at day 22, followed by Dunnett’s multiple comparisons test (**f**).

