## Supplemental Methods for "Inhibition of the IL-17A axis Protects against Immune-related Adverse Events while Supporting Checkpoint Inhibitor Anti-tumor Efficacy"

**Lechner *et al.* Supplemental Methods**

**Anti-mouse antibodies used in flow cytometry experiments.**

| Target | clone | Source | Working Dilution | Fluorophore | Catalog number |
|---|---|---|---|---|---|
| CD4 | RM4-5 | Invitrogen | 1:50 | FITC | 11-0042-85 |
| CD3 | 145-2C11 | Invitrogen | 1:50 | APC | 17-0031-82 |
| CD8 | 53-6.7 | Invitrogen | 1:50 | APC | 17-0081-82 |
| RORgt | AFKJS-9 | eBioscience | 1:50 | APC | 17-6988-82 |
| IL-17A | eBio17B7 | Invitrogen | 1:50 | PE | 12-7177-81 |
| CD11c | N418 | Invitrogen | 1:50 | PE | 12-0114-82 |
| CD11b | M1/70 | Invitrogen | 1:50 | APC | 17-0112-82 |
| F4/80 | BM8 | Invitrogen | 1:50 | PE | 12-4801-80 |
| B220 | RA3-6B2 | eBioscience | 1:50 | e450 | 48-0452-82 |
| Nkp46 | 29A1.4 | Invitrogen | 1:50 | PeCy7 | 25-3351-82 |
| TCRgd | GL3 | eBioscience | 1:50 | e450 | 48-5711-82 |
| CD45 | 30-F11 | Biolegend | 1:50 | PerCP Cy5.5 | 103132 |
| CD45 | 30-F11 | Invitrogen | 1:50 | SB600 | 63-0451-82 |
| CD45 | 30-F11 | Invitrogen | 1:50 | PE | 12-0451-82 |

**Anti-human antibodies used in flow cytometry experiments.**

| Target | clone | Source | Working Dilution | Fluorophore | Catalog number |
|---|---|---|---|---|---|
| CD45 | H130 | Biolegend | 1:50 | PE | 304008 |
| CD3 | SK7 | Invitrogen | 1:50 | PeCy7 | 25-0036-42 |
| CD19 | 4G7 | Biolegend | 1:50 | APC | 392504 |
| CD14 | 61D3 | Invitrogen | 1:50 | FITC | 11-0149-42 |

**Taqman probes used in qRT-PCR experiments.**

| Gene | Species | Probe (Assay ID) | Source |
|---|---|---|---|
| GAPDH | mouse | Mm99999915_g1 | Thermo Fisher |
| IL-17A | mouse | Mm00439618_m1 | Thermo Fisher |
| IL-1b | mouse | Mm00434228_m1 | Thermo Fisher |
| IL-6 | mouse | Mm00446190_m1 | Thermo Fisher |
| TNFa | mouse | Mm00443258_m1 | Thermo Fisher |
| Tgfb1 | mouse | Mm01178820_m1 | Thermo Fisher |
| RORC | mouse | Mm01261022_m1 | Thermo Fisher |
| IL23R | mouse | Mm00519943_m1 | Thermo Fisher |
| IFNG | mouse | Mm01168134_m1 | Thermo Fisher |
| CD3e | mouse | Mm00599683_m1 | Thermo Fisher |
| ptprc | mouse | Mm01293577_m1 | Thermo Fisher |
