## Extended Data Table 1 for "Inhibition of the IL-17A axis Protects against Immune-related Adverse Events while Supporting Checkpoint Inhibitor Anti-tumor Efficacy"

**ICI-Thyroiditis Patients**

| Flow cytometry sample | scRNAseq sample | Diagnosis | Sex | Age (Years) | Cancer type | Immunotherapy | Time to IrAE (weeks) | TPO Ab | Tg Ab | LT4 therapy | Best response | Other IrAEs |
|---|---|---|---|---|---|---|---|---|---|---|---|---|
| x | | Thyroid IrAE | Male | 85.3 | Melanoma | Nivolumab | 11.7 | - | - | Y | Stable disease | |
| x | x | Thyroid IrAE | Female | 30.0 | Leiomyosarcoma | Ipilimumab + Nivolumab | 6.0 | - | + | Y | Progression of disease | |
| | x | Thyroid IrAE | Male | 57.3 | Melanoma | Ipilimumab + Nivolumab | 9.0 | - | - | Y | Partial response | |
| | x | Thyroid IrAE | Male | 58.0 | Sarcoma | Pembrolizumab | 3.0 | - | - | Y | Progression of disease | |
| x | x | Thyroid IrAE | Male | 59.3 | Non-small cell lung cancer | Durvalumab | 4.0 | - | - | Y | Stable disease | |
| x | | Thyroid IrAE | Male | 47.6 | Renal cell carcinoma | Pembrolizumab | 9.0 | - | - | Y | Stable disease | Hypophysitis |
| x | | Thyroid IrAE | Male | 66.0 | Hepatocellular carcinoma | Pembrolizumab | 10.6 | + | - | Y | Stable disease | |
| x | | Thyroid IrAE | Female | 46.3 | High grade neuroendocrine tumor | Atezolizumab | 18.7 | - | + | Y | Progression of disease | |

**Hashimoto's thyroiditis Patients**

| Flow cytometry sample | scRNAseq sample | Diagnosis | Sex | Age (Years) | | | | TPO Ab | Tg Ab | LT4 therapy |
|---|---|---|---|---|---|---|---|---|---|---|
| x | x | HT | Male | 74.6 | | | | + | + | Y |
| x | x | HT | Female | 35.1 | | | | + | - | Y |
| x | x | HT | Female | 51.6 | | | | + | ND | Y |
| x | | HT | Female | 28.5 | | | | + | - | Y |
| x | | HT | Female | 57.3 | | | | + | + | N |
| x | | HT | Female | 53.0 | | | | + | ND | Y |
