## Extended Data Figure 1 for "Inhibition of the IL-17A axis Protects against Immune-related Adverse Events while Supporting Checkpoint Inhibitor Anti-tumor Efficacy"

**Lechner et al. Extended. Data Fig. 1**

**a Gating strategy for immune cell populations in patient thyroid fine needle aspiration (FNA) specimens**

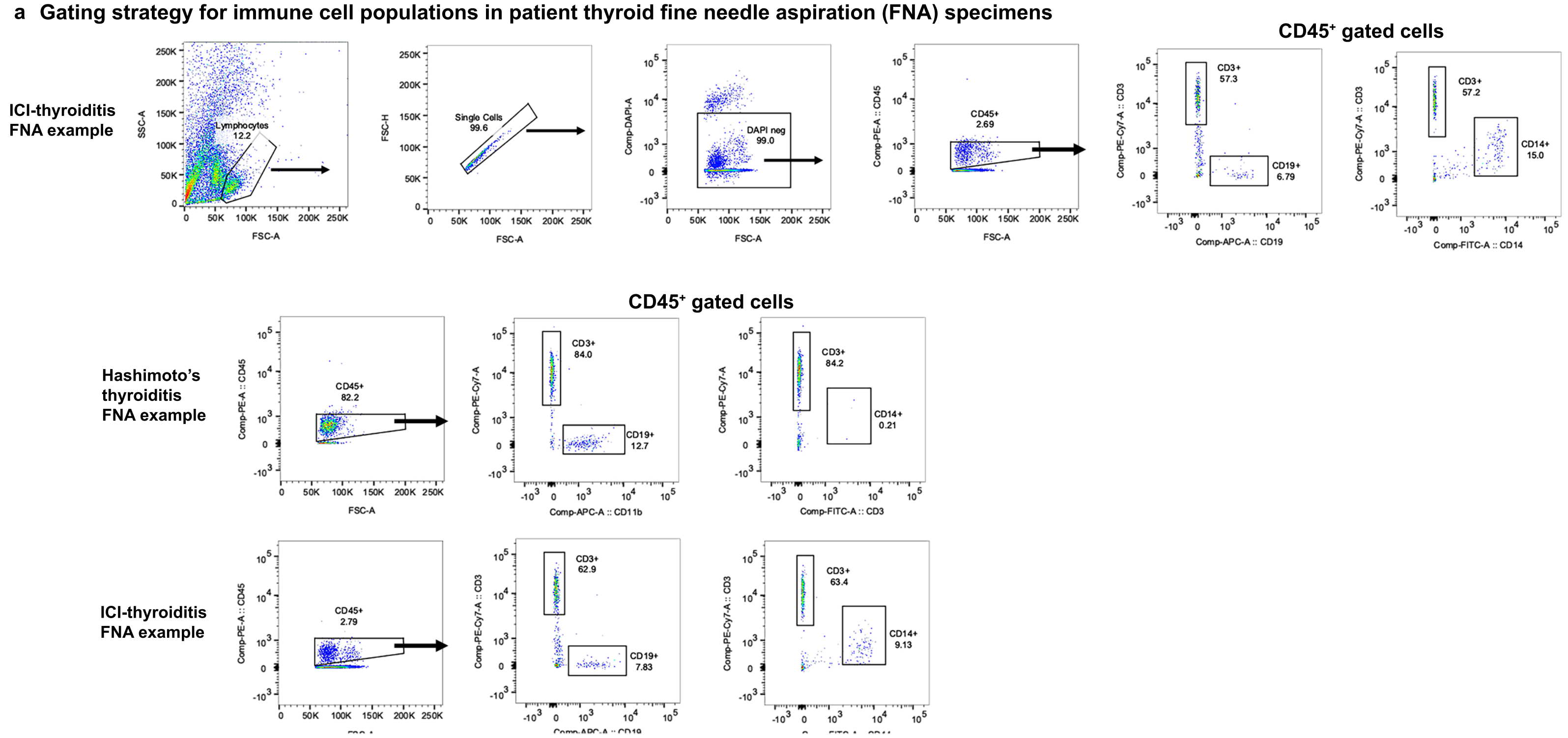


**b Gene expression across cell clusters of human immune cells from thyroid FNA specumens by scRNAseq**

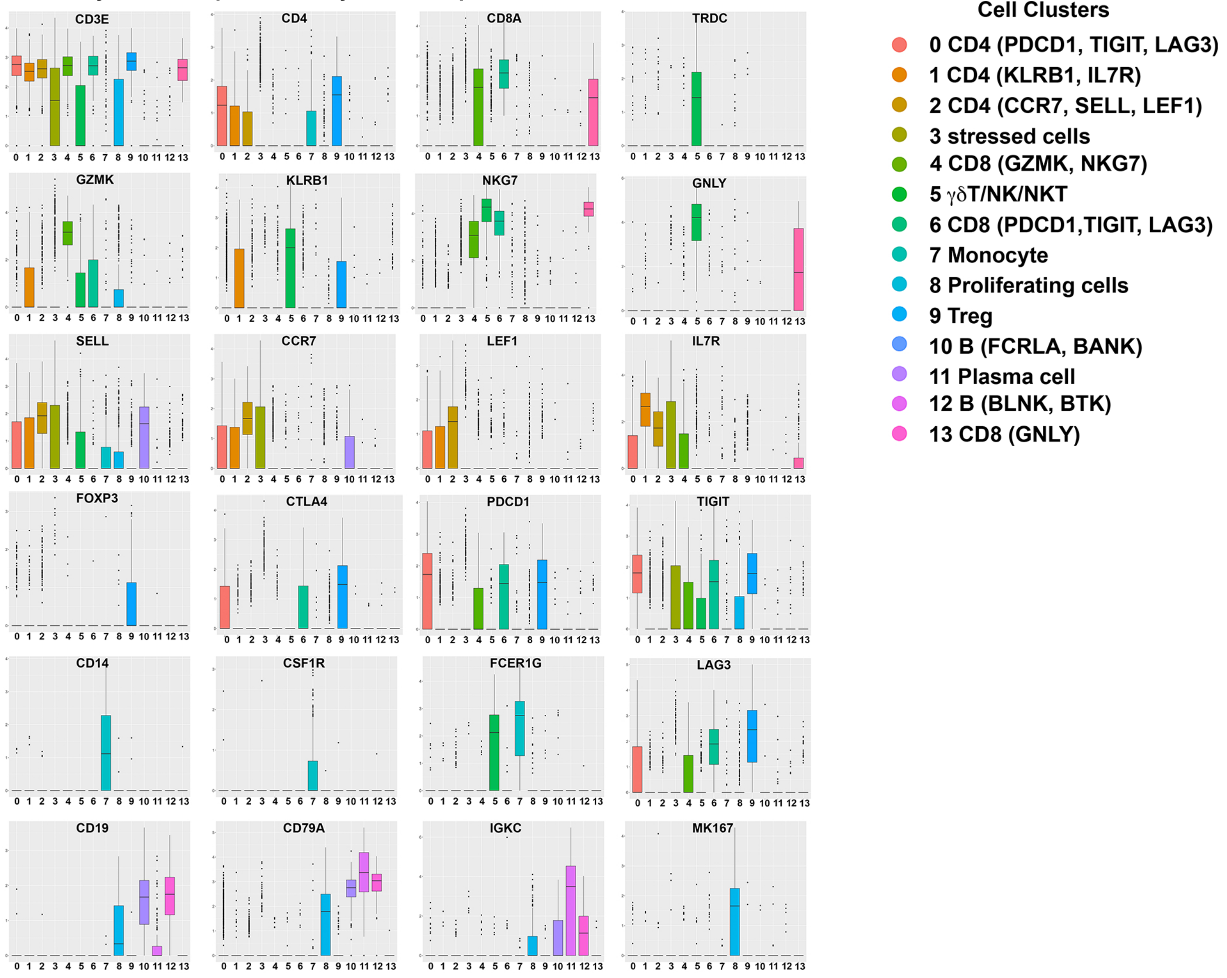


**Extended Data Fig. 1. Supplemental data for flow cytometry and scRNAseq of human thyroid fine needle aspiration (FNA) specimens from Hashimoto's thyroiditis (HT) and immune checkpoint inhibitor (ICI)-thyroiditis patients. a,** Gating strategy and representative dot plots for immune cells. **b**, The expression levels of various marker genes of each cluster in human thyroid specimen UMAP clusters for relevant immune populations
