## Extended Data Figure 2 for "Inhibition of the IL-17A axis Protects against Immune-related Adverse Events while Supporting Checkpoint Inhibitor Anti-tumor Efficacy"

Lechner et al. Extended Data Fig. 2

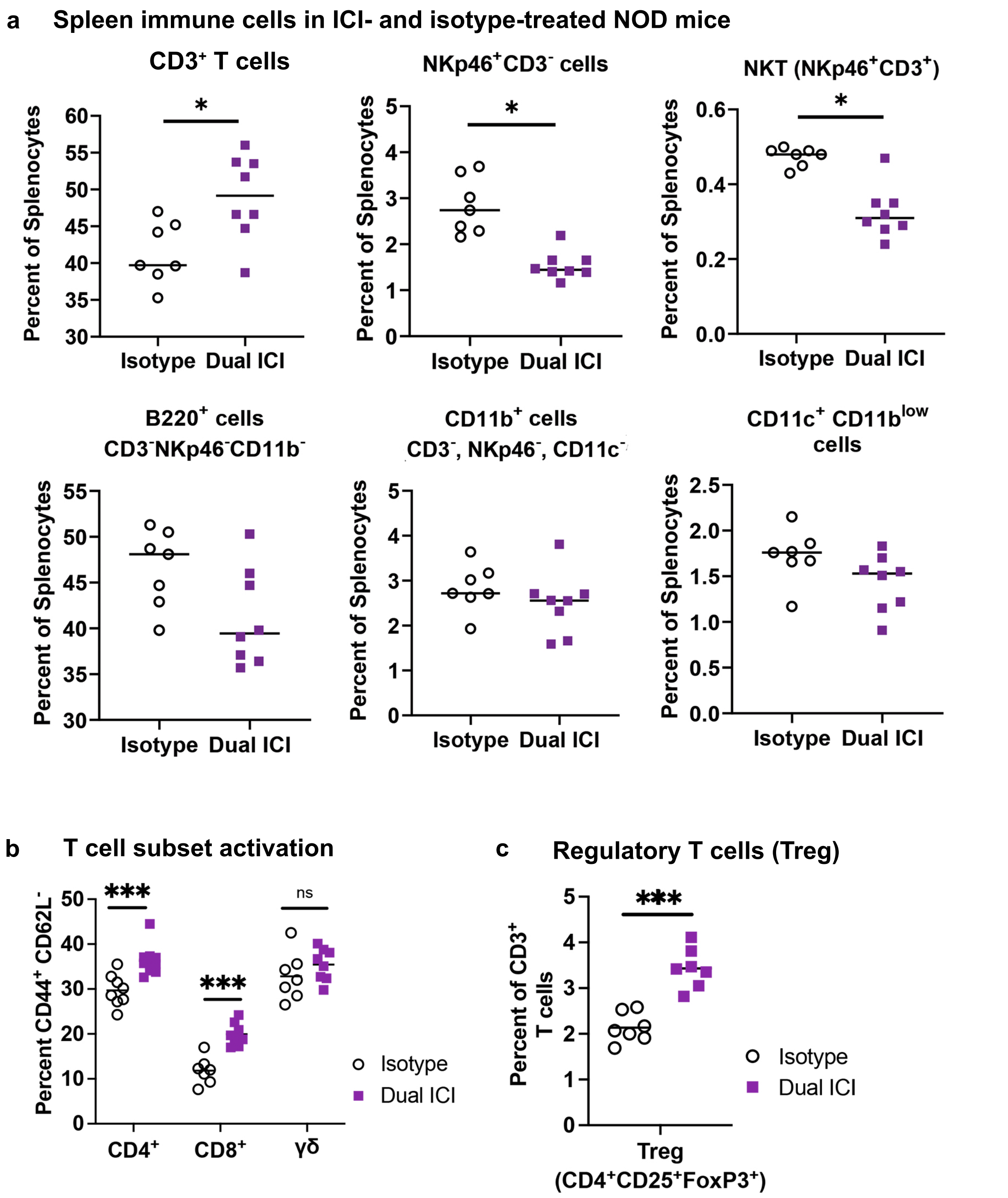


**Extended Data Fig. 2. Flow cytometry analysis of peripheral immune changes in ICI-treated NOD mice**. **a,** Relative frequency of putative T ($CD3^+$ $NKp46^-$), natural killer (NK, $NKp46^+$ $CD3^-$), NKT ($CD3^+$ $NKp46^+$), B ($B220^+$ $CD3^-$ $NKp46^-$ $CD11b^-$), myeloid ($CD11b^+$) and conventional dendritic cell (DC, $CD11c^+$ $CD11b^{low}$) as a percent of splenocytes in isotype (*n*=7) *vs.* anti-PD-1 + anti-CTLA-4 (Dual ICI, *n*=8) mice after 4 weeks of treatment. P values are as follows: T cell 0.012, NK cell 0.004, NKT cell 0.002, B cell 0.12, $CD11b^+$ myeloid cell 0.3, and DC 0.16. **b,** Percent of $CD4^+$, $CD8^+$ and $\gamma\delta TCR^+$ ($\gamma\delta$) T cells in spleens of Dual ICI-treated (*n*=8) or isotype (*n*=8) mice expressing activation markers ($CD44^+$ $CD62L^-$), by flow cytometry. P values are as follows: $CD4^+$ 0.002, $CD8^+$ 0.0003, $\gamma\delta$ 0.29. d, Relative frequency of regulatory T cells within CD3+ cells in spleen of ICI-treated (*n*=8) or isotype (*n*=7) mice, p=0.0001. Data shown are mean ± SEM. *p<0.05, **p<0.01, ***p<0.001. Differences in immune populations in spleen were compared by two-tailed, unpaired t test with Welch correction, assuming unequal s.d., and Holm-Sidak method correction for multiple comparisons.
