## Extended Data Figure 3 for "Inhibition of the IL-17A axis Protects against Immune-related Adverse Events while Supporting Checkpoint Inhibitor Anti-tumor Efficacy"

### Lechner et al. Extended Data Fig. 3

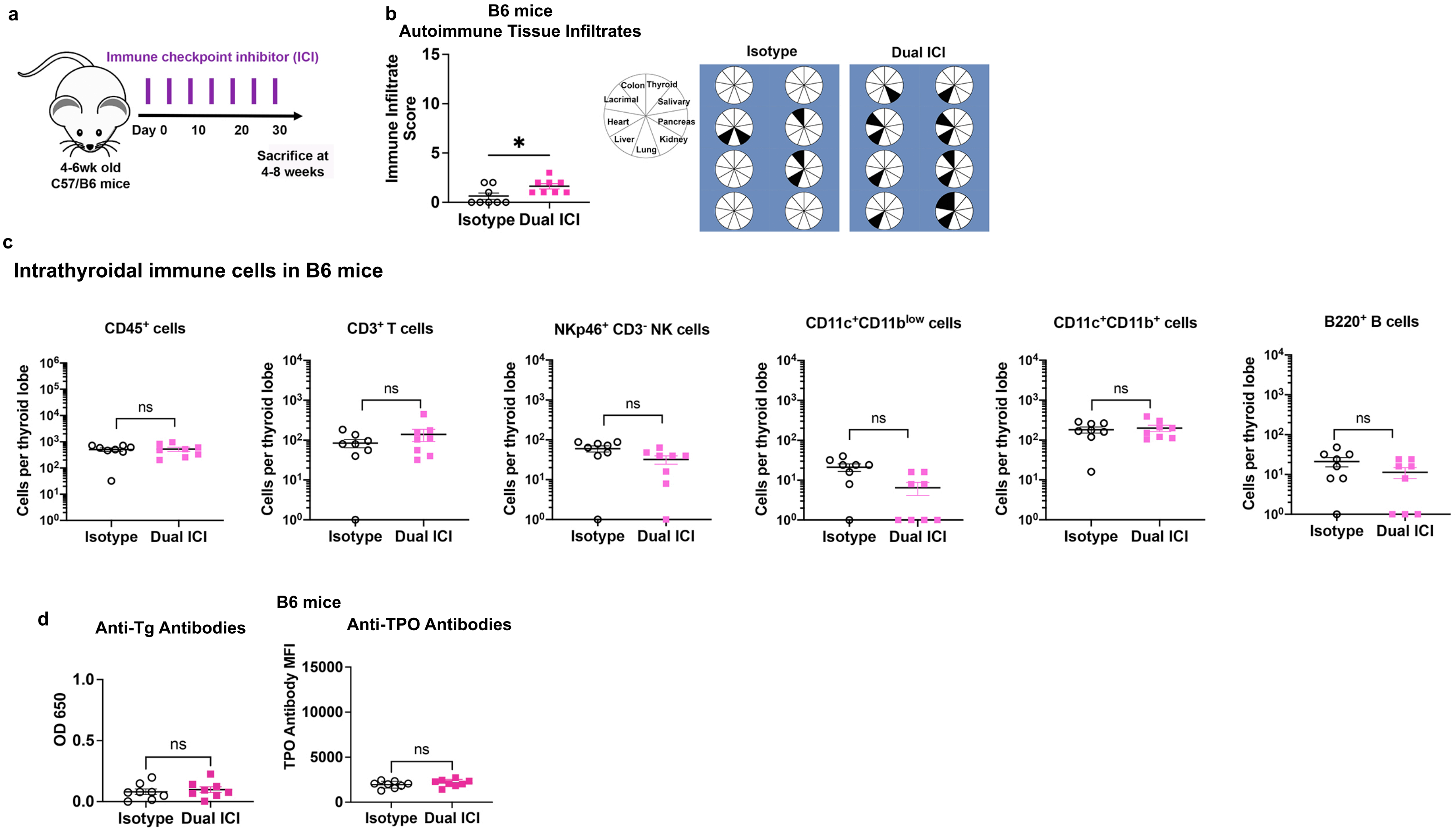


**Extended Data Fig. 3. ICI therapy in B6 mice induces minimal tissue autoimmunity.**
**a**, Schematic of ICI drug treatment. **b**, Comparison of autoimmune organ infiltration after 4 weeks of ICI *vs.* isotype treatment in C57/B6 (B6) mice ($n$=8 isotype, $n$=8 Dual ICI); Pie charts (*right*) show tissue infiltrate for each mouse; black = immune infiltrate, white = no infiltrate. **c,** Flow cytometry analysis of thyroid-infiltrating immune cells in anti-PD-1 + anti-CTLA-4 (Dual ICI) treated B6 mice. Mice were euthanized after 4 weeks of ICI or isotype treatment, perfused with saline, and then fresh thyroid tissues dissociated into single cell suspensions. Cells were stained with fluorescent antibodies and analyzed by flow cytometry, with estimation of cells/thyroid lobe for each animal. Each point represents an individual mouse ($n$=8 isotype and $n$=8 Dual ICI). **d**, Quantification of anti-thyroid autoantibodies in B6 mice after 4 weeks of isotype ($n$=8) or Dual ICI ($n$=8) treatment. Data are mean±SEM. Groups were compared by two-tailed, unpaired t test with Welch correction, assuming unequal s.d., and Holm-Sidak method correction for multiple comparisons (**b, c, d**). *p<0.05.
