## Extended Data Figure 4 for "Inhibition of the IL-17A axis Protects against Immune-related Adverse Events while Supporting Checkpoint Inhibitor Anti-tumor Efficacy"

### Lechner et al. Extended Data Fig. 4

**a** Gene expression across cell clusters in mouse intrathyroidal immune cells by scRNAseq

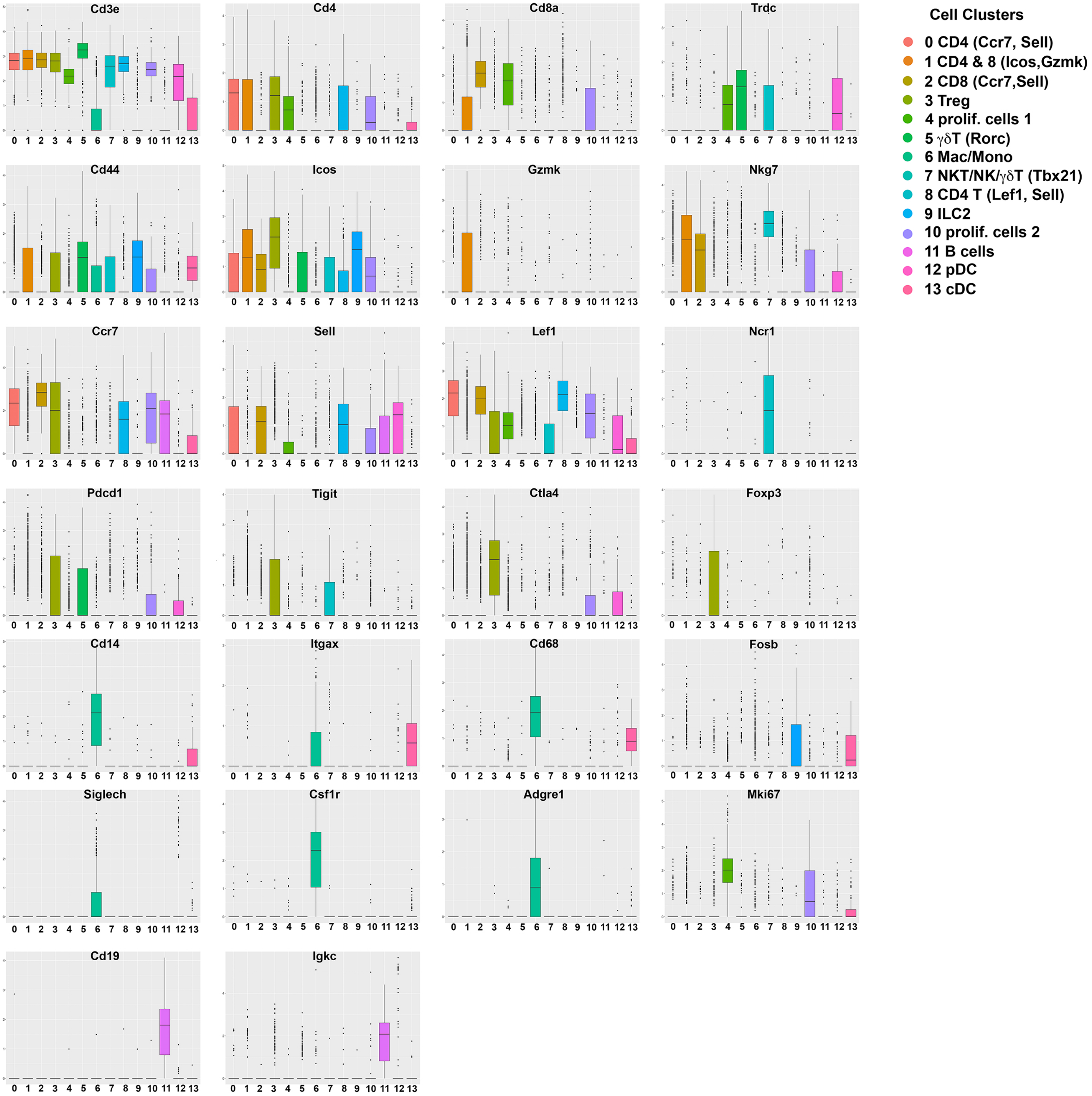


**b** Type 3 immune response

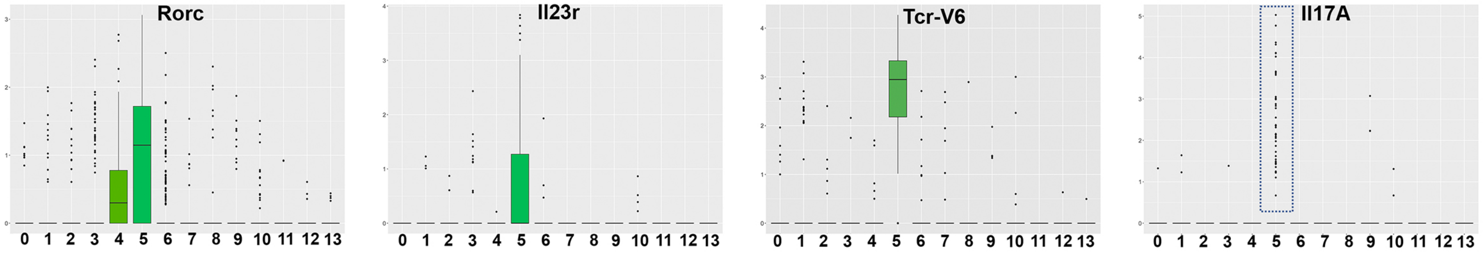


**Extended Data Fig. 4. Supplemental data for scRNAseq of thyroid immune infiltrates from isotype or Dual ICI-treated NOD mice**. **a**, The expression levels of various marker genes of each cluster in mouse thyroid tissue UMAP clusters for relevant immune populations. **b,** Expression of immune genes associated with Type 3 immune responses across cell clusters.
