## Extended Data Figure 5 for "Inhibition of the IL-17A axis Protects against Immune-related Adverse Events while Supporting Checkpoint Inhibitor Anti-tumor Efficacy"

### Lechner et al. Extended Data Fig. 5

#### Gating strategy for mouse primary immune cells

**a** Example of gating for immune populations (spleen control shown)

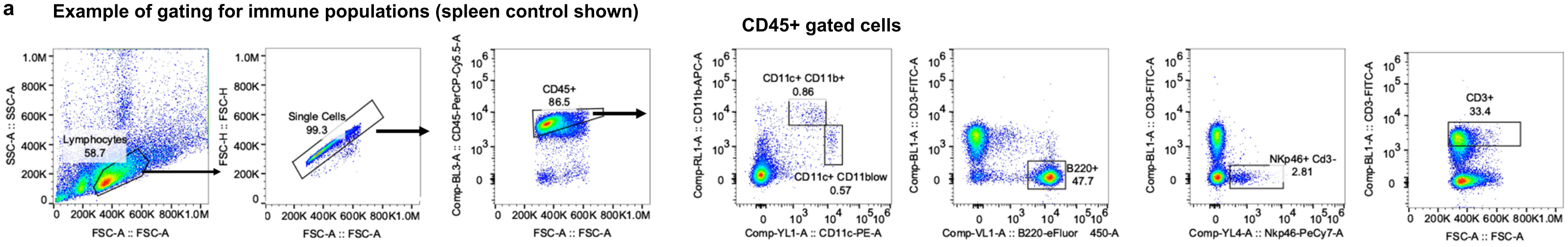


**b** Example of CD45+ and T cell gating (thyroid specimen shown)

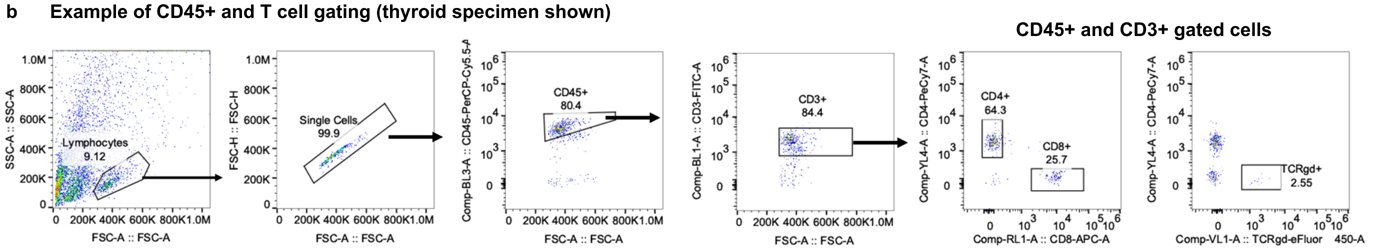


**c** Example of IL-17A+ staining in thyroid specimens showing CD3+ and CD3- populations

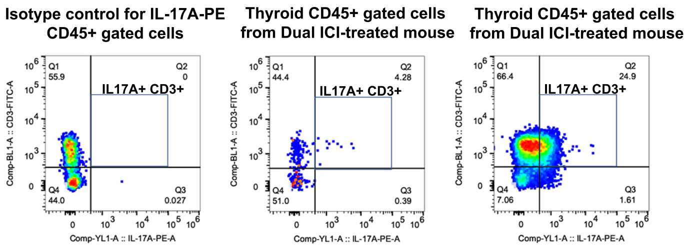


**Extended Data Fig. 5. Gating strategy and representative dot plots for primary mouse immune cells. a,** Gating strategy and representative dot plots for CD45$^+$, CD3$^+$, Nkp46$^+$, B220$^+$, CD11c$^+$ CD11b$^+$, and CD11c$^+$ CD11b$^{low}$ cells. **b,** Gating strategy and representative dot plots for CD4$^+$, CD8$^+$, and TCRγδ$^+$ T cell subsets. **c,** Representative dot plots for IL-17A$^+$ staining of CD45$^+$ immune cells showing CD3$^+$ and CD3$^-$ populations, and variability in the amount of intrathyroidal immune cells across ICI-treated specimens (*center* and *right* panels).
