## Extended Data Figure 6 for "Inhibition of the IL-17A axis Protects against Immune-related Adverse Events while Supporting Checkpoint Inhibitor Anti-tumor Efficacy"

Lechner et al. Extended Data Fig. 6

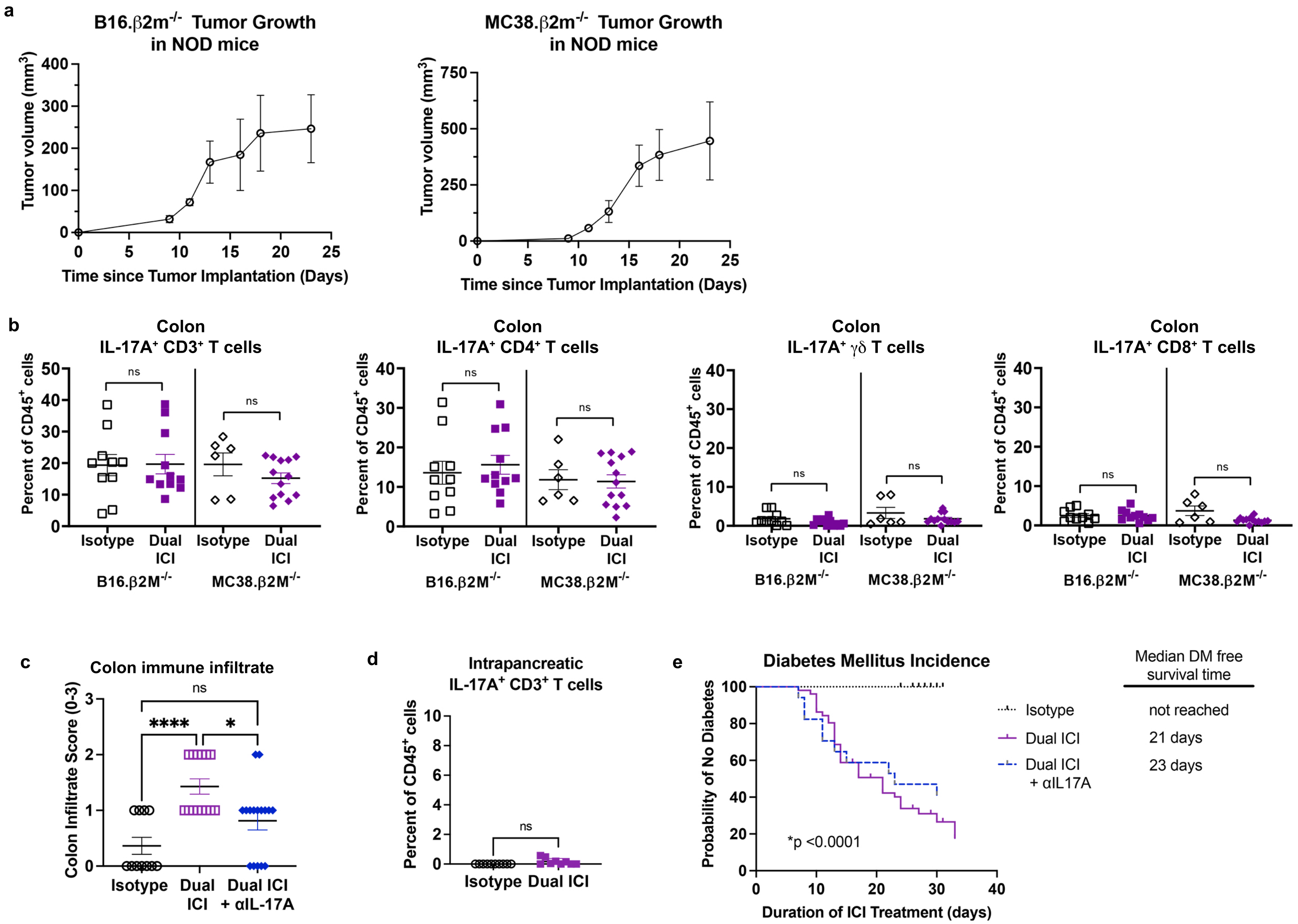


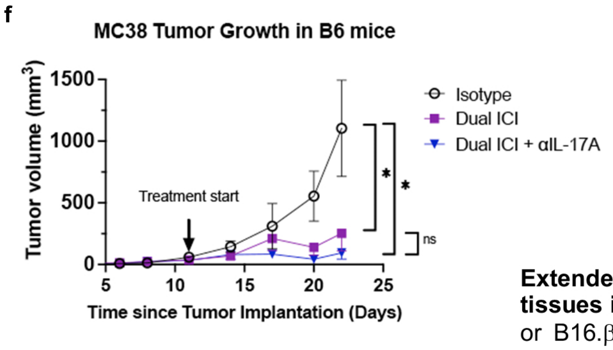


**Extended Data Fig. 6. Tumor growth and ICI-associated IL-17A$^+$ T cells in non-thyroid tissues in NOD mice. a,** Representative growth curves for tumors in NOD mice inoculated with or B16.$\beta 2M^{-/-}$ melanoma or MC38.$\beta 2M^{-/-}$ colon carcinoma tumor cells. **b,** Relative frequency among CD45$^+$ immune cells of IL-17A$^+$ CD3$^+$ T cells and IL-17A$^+$ T cell subsets isolated from colon tissue of isotype or Dual ICI-treated mice at 4 weeks by flow cytometry, 3 experiments, each dot represents a single animal. **c***,* Comparison of colon immune infiltrate on histology for isotype (*n*=11), Dual ICI-treated (*n*=11), or Dual ICI and neutralizing IL-17A ($\alpha$IL17A) therapy (*n*=14) NOD mice after 4 weeks, 2 experiments, each dot represents a single animal. **d**, Relative frequency among CD45$^+$ immune cells of IL-17A$^+$ T cells in the pancreas of isotype *vs.* Dual ICI-treated mice, 2 experiments, each dot represents a single animal. **e,** Kaplan-Meier curve showing incident autoimmune diabetes mellitus in NOD mice among treatment groups as assessed by persistent glucosuria requiring insulin therapy. **f,** Growth of syngeneic MC38 colon tumors in C57/B6 (B6) mice treated with isotype, Dual ICI, or Dual ICI and $\alpha$IL17A, *n*=6-8 each group. Data are mean $\pm$SEM shown. *p<0.05, **p<0.01, ***p<0.001, ****p<0.0001. Brown-Forsythe ANOVA, assuming unequal s.d., followed by Dunnett's multiple comparisons test (**b,c**); Two-tailed, unpaired t test with Welch correction, assuming unequal s.d. and Holm-Sidak method correction for multiple comparisons (**d**); Log rank test for trend for median diabetes-free survival (**e**); or ANOVA for tumor volume at day 22, followed by Dunnett's multiple comparisons test (**f**).
